# Secretory MPP3 reinforce myeloid differentiation trajectory and amplify myeloid cell production

**DOI:** 10.1101/2021.09.01.458573

**Authors:** Yoon-A Kang, Hyojung Paik, Si Yi Zhang, Jonathan Chen, Matthew R. Warr, Rong Fan, Emmanuelle Passegué

**Author notes:** Corresponding and lead author: Emmanuelle Passegué, PhD, Alumni Professor of Genetics & Development, Director of the Columbia Stem Cell Initiative, Columbia University Irving Medical Center, 650 West 168^th^ Street, BB-1109A, New York, NY 10032, USA. These authors contributed equally to this work (Yoon-A Kang, Hyojung Paik).

## Abstract

Recent lineage tracing analyses revealed multipotent progenitors (MPP) to be major functional contributors to steady-state hematopoiesis (*1–6*). However, we are still lacking a precise resolution of myeloid differentiation trajectories and cellular heterogeneity in MPPs. Here, we found that myeloid-biased MPP3 (*2, 3*) are functionally and molecularly heterogeneous, with a distinct subset of myeloid-primed secretory cells with high endoplasmic reticulum (ER) volume and Fc*γ*R expression. We show that Fc*γ*R^+^/ER^high^ MPP3 are a transitional population for rapid production of granulocyte/macrophage progenitors (GMP), which directly amplify myelopoiesis through inflammation-triggered secretion of cytokines in the local bone marrow (BM) microenvironment. Our results identify a novel regulatory function for a subset of secretory MPP3 that controls myeloid differentiation through lineage-priming and cytokine production, and act as a self-reinforcing amplification compartment in stress and disease conditions.

**One-Sentence Summary:** A secretory subset of multipotent hematopoietic progenitors augment myelopoiesis in stress and diseases conditions.

## Main Text

Myelopoiesis is a demand-adapted process where hematopoietic stem cells (HSC) and a collection of progenitor cells integrate signals from their environment and tailor the output of the myeloid lineage to meet the specific needs of the organism and respond to physiological challenges (*1, 7, 8*). Emergency myelopoiesis is induced to amplify myeloid cell production either acutely in stress conditions or constitutively in various malignant contexts, and resulting in a major re- organization of the hematopoietic stem and progenitor cell (HSPC) compartment at the top of the hematopoietic hierarchy (*1, 9*). Quiescent HSCs are first activated leading to expansion of myeloid-biased MPP2 and MPP3 and myeloid-reprogramming of lymphoid-biased MPP4, which result in the formation of self-renewing GMP patches and their expansion into GMP clusters driving local burst production of mature myeloid cells in the BM microenvironment (*2, 10, 11*). The remodeling of the MPP compartment is triggered in part by low Notch and high Wnt activity in HSCs, and by pro-inflammatory cytokines like IL-1, TNF*α*, or IL-6 that stimulate many steps of this regenerative program to amplify myeloid cell production (*10–14*). Interestingly, HSPCs themselves have been shown to secrete cytokines upon inflammatory stimuli (*15*), raising the intriguing possibility that autocrine or paracrine signaling in the local BM niche might control their function. Single cell RNA-sequencing (scRNAseq) analyses and barcoding lineage tracing strategies have also considerably advanced our knowledge of lineage specification and revealed molecular and functional heterogeneity as well as mixed lineage expression patterns in distinct HSPC populations (*16–19*). However, we still lack a clear understanding of the regulatory mechanisms enforcing myeloid lineage commitment at the top of the hematopoietic hierarchy, and the precise role of lineage-biased MPPs in amplifying myeloid cell production. With the current SARS-CoV-2 pandemic and the success of immunotherapy for cancer treatment, there is a well-justified interest in understanding the mechanisms controlling immune cell production particularly as it relates to innate immunity and myeloid cell production. Indeed, studies of “trained immunity” and the discovery of “central trained immunity” have demonstrated the importance of myeloid progenitors in the regulation of innate immune responses (*20*). Here, we show that a subset of secretory MPP3 acts as a self-reinforcing amplification compartment controlling myeloid differentiation in stress and disease conditions through both lineage-priming and differential cytokine production in the local BM niche microenvironment.

### Identification of a secretory subset of MPP3

To further characterize HSPC secretory activity, we focused on several well-defined phenotypic populations including HSCs (Lin^-^/c-Kit^+^/Sca-1^+^/Flk2^-^/CD48^-^/CD150^+^), myeloid-biased MPP3 (Lin^-^/c-Kit^+^/Sca-1^+^/Flk2^-^/CD48^+^/CD150^-^) and lymphoid-biased MPP4 (Lin^-^/c-Kit^+^/Sca-1^+^/Flk2^+^), which we compared to myeloid-committed GMPs (Lin^-^/c-Kit^+^/Sca-1^-^/Fc*γ*R^+^/CD34^+^) (fig. S1A). We found that ∼ 1/3 of MPP3 had high endoplasmic reticulum (ER) volume by transmission electron microscopy (TEM) and confirmed the presence of an ER^high^ subset of MPP3 by immunofluorescence microscopy for the ER marker KDEL and flow cytometry staining for the ER-Tracker dye (Fig. 1, A and B, and fig. S1, B and C). Interestingly, the rough ER structure observed in MPP3 was morphologically similar to the secretory apparatus found in specialized immunoglobulin-producing plasma cells, although MPP3 shared no surface markers with plasma cells (fig. S1, C and D). We directly tested the secretory activity of MPP3 by treating isolated HSPC populations with a previously described inflammatory LPS/Pam3CSK4 (L/P) stimulus (*15*), and collecting supernatants after 24 hours for secretome analyses. Stimulated MPP3 secreted higher levels of TNF*α* and IL-6 than HSCs and MPP4 as shown by ELISA, and displayed a global increase in cytokine secretion akin to GMP as measured with the Raybiotech 200 mouse cytokine array (Fig. 1, C and D). Detailed examination of each HSPC population revealed specific and complex secretory patterns, with stimulated MPP3 showing increased secretion of many pro-inflammatory and pro-myeloid differentiation cytokines including IL-1*α*, G-CSF and GM-CSF, and decreased production of regulatory factors controlling immune cell function like TACI or CD40L (Fig. 1E, fig. S1E, and table S1). Consistently, MPP3 also expressed unfolded protein response (UPR) genes in the range of GMPs but to a lower extent than plasma cells (fig. S1F). We next investigated MPP3 secretion at the single cell level using 14 pre-selected cytokines and an established nanofluidic technology (*21*) (fig. S2A). We confirmed the higher overall secretory activity in MPP3 compared to HSCs and MPP4 (fig. S2B). However, at the single cell level, stimulated MPP3 secreted less TNF*α* and IL-6 than unstimulated MPP3, which was likely due to their concomitant production of IL-10, a known suppressor of IL-6 and TNF*α* secretion (*22*) (fig.

**Fig. 1.**
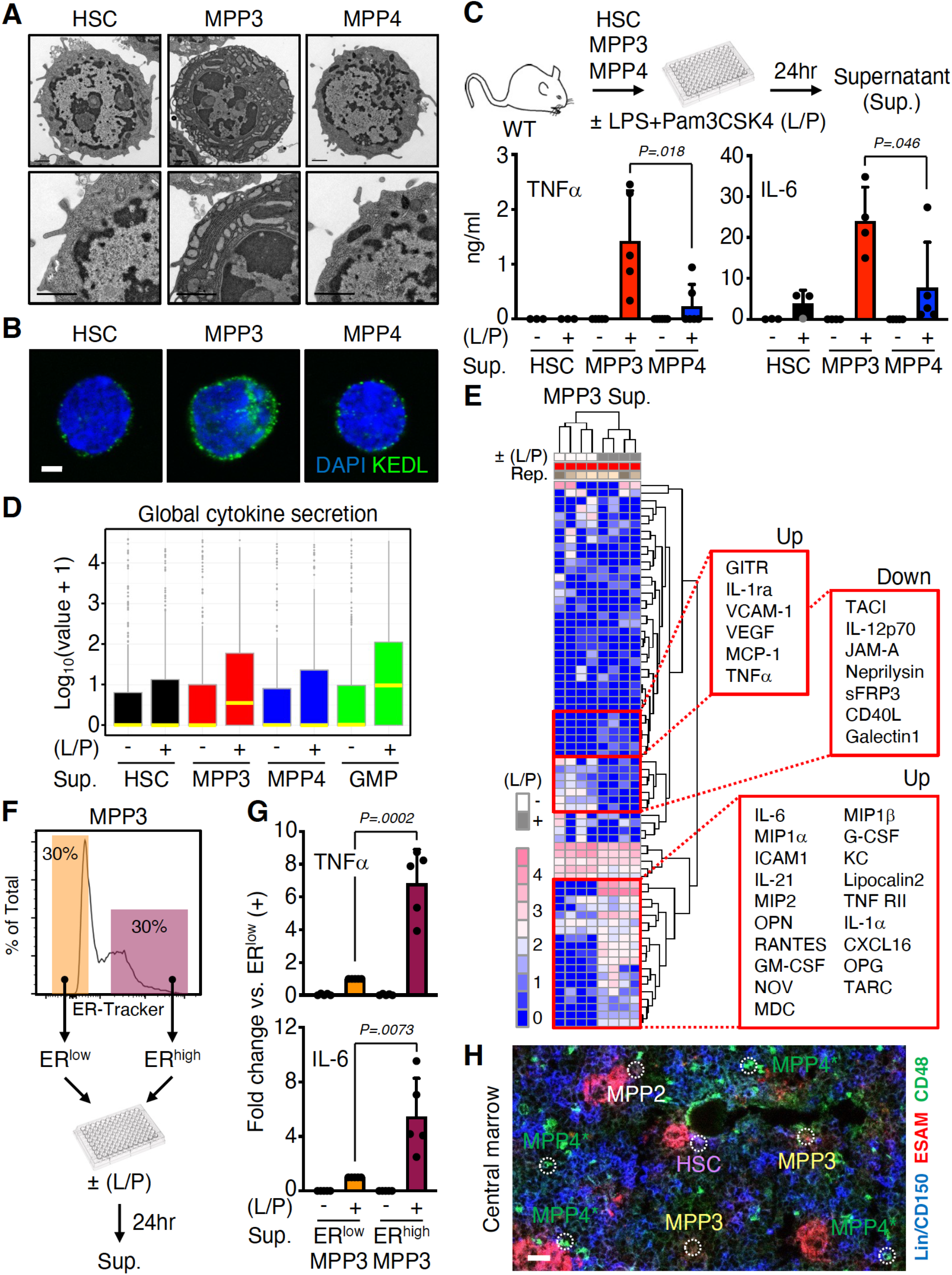
Secretory features of MPP3. (**A**) Representative transmission electron microscopy images of HSC, MPP3 and MPP4. Scale bar, 1 μm. (**B**) Representative immunofluorescence images of HSC, MPP3 and MPP4 stained for the endoplasmic reticulum (ER) marker KEDL. Scale bar, 3 μm. (**C**) Differential secretion of TNF*α*and IL-6 by HSC, MPP3 and MPP4 upon stimulation. Experimental scheme and results from ELISA measurements are shown. Supernatants were collected upon culture of 10,000 cells for 24 hours in 150 µl base media with or without (±) LPS/Pam3CSK4 (L/P) stimulation. (**D** and **E**) Stimulated MPP3 are the most secretory HSPCs with: (D) box plots of secreted cytokine intensity by HSC, MPP3 and MPP4 upon stimulation (yellow line, mean value), and (E) heatmap of unsupervised clustering of MPP3-secreted cytokines after quantile normalization (representative increased and decreased cytokines upon stimulation are indicated on the right). Results are from 24 hours supernatants analyzed with the Raybiotech 200 mouse cytokine array. (**F**) Experimental scheme for isolating and analyzing ER^high^ (top 30% of ER-Tracker staining) and ER^low^ (bottom 30% of ER-Tracker staining) MPP3 subsets. (**G**) Differential secretion of IL-6 and TNF*α* by ER^high^ vs. ER^low^ MPP3 upon stimulation. Results are from 24 hours supernatants analyzed by Luminex cytokine bead array. (**H**) Representative image of *in situ* immunofluorescence staining of HSPCs in the central marrow cavity. HSC, MPP2, MPP3 and MPP4/GMP are indicated by white dotted line circles. Asterisk (*) denotes the marker overlap between MPP4 and GMPs with this staining scheme. Scale bar, 20 μm. Data are means ± S.D. except when indicated, and significance was assessed by a two-tailed unpaired Student’s t-test.

S2, C to E). This negative self-regulatory effect of IL-10 was likely overcome in bulk culture due to the strong pro-secretion effect of other cytokines including TNF*α*itself, which directly modulated IL-6 secretion (fig. S2F). We also confirmed a role for the classical mechanisms regulating cellular secretion (*23, 24*), with inhibition of NF-*κ*B and Ca^2+^-dependent signaling impairing IL-6 secretion from stimulated MPP3 (fig. S2F). To directly separate high from low secretory MPP3, we took advantage of their difference in ER volume and sub-fractionated MPP3 into ER^high^ (top 30%) and ER^low^ (bottom 30%) subsets based on ER-Tracker staining (Fig. 1F). We confirmed significantly higher secretion of many cytokines, including TNF*α* and IL-6, in stimulated ER^high^ MPP3 (Fig. 1G and fig. S2G). To provide further *in vivo* support to the paracrine effect of MPP3 secretion, we developed a novel immunofluorescence imaging panel to visualize MPP3 in their BM niche microenvironment (fig. S3, A and B). Imaging BM sections confirmed the close proximity of MPP3 with other HSPC populations including HSCs, MPP4 and GMPs (Fig. 1H and fig. S3C). Taken together, these results demonstrate that MPP3 are the most secretory HSPCs, identify a unique subset of ER^high^ MPP3 that robustly produce many pro-inflammatory/myeloid differentiation cytokines upon inflammatory stimulus, and localize MPP3 in the vicinity of other HSPCs in the BM niche in support of a local paracrine and autocrine effect of MPP3 secretion in regulating myelopoiesis.

### Molecular heterogeneity in MPP3

To gain a better understanding of MPP3 heterogeneity and response to inflammatory stimulation, we performed droplet-based single cell RNA-sequencing (scRNAseq) analyses on MPP3 stimulated with or without LPS/Pam3CSK4 for 6 hours. Data from unstimulated and stimulated MPP3 were harmonized by nearest neighbor integration (*25*) and uniform manifold approximation and projection (UMAP) representation identified 13 different clusters (Fig. 2A).

**Fig. 2.**
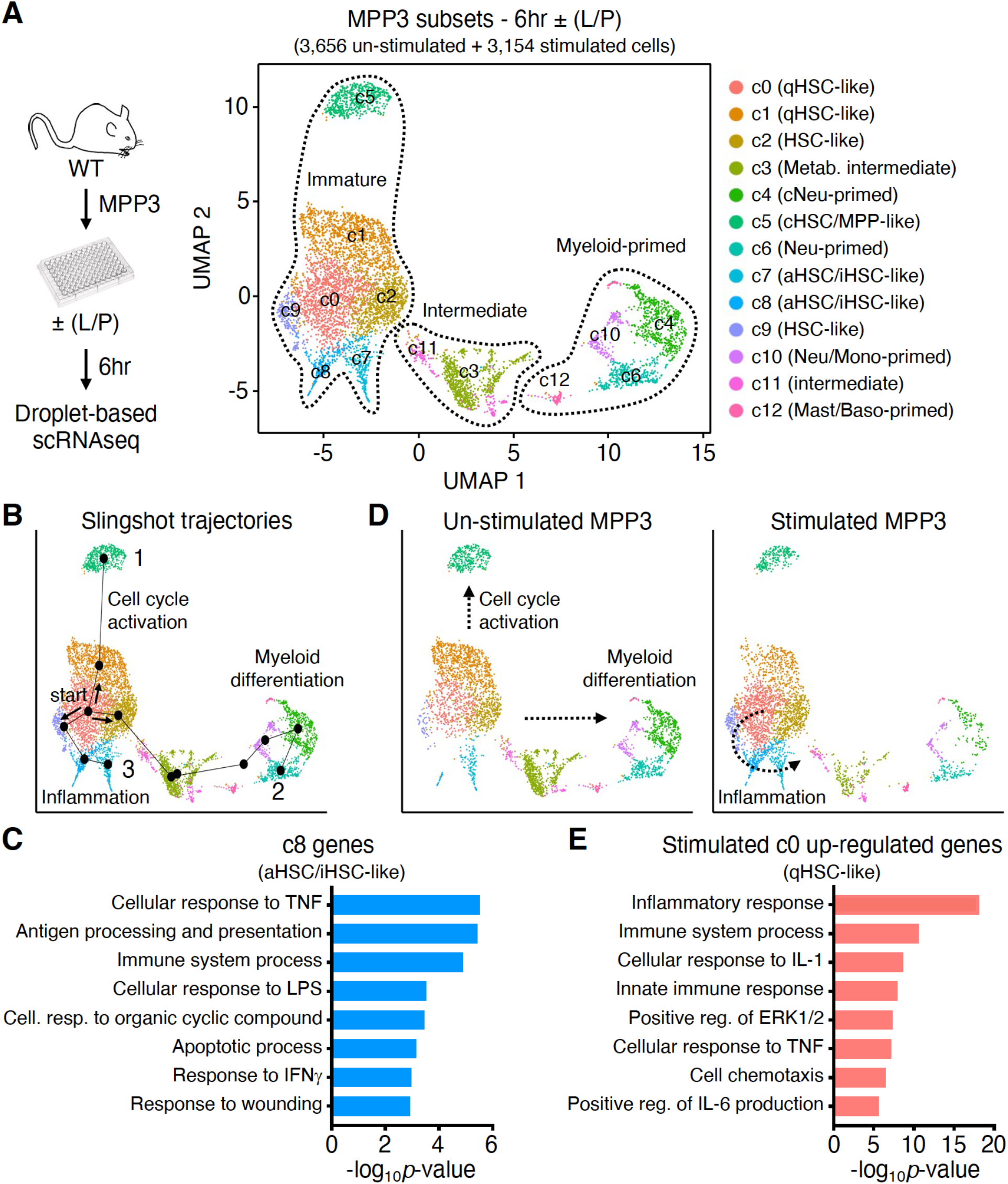
MPP3 heterogeneity and inflammatory remodeling. (**A**) UMAP representation of un- stimulated/stimulated MPP3 scRNAseq dataset with experimental scheme and cluster identification; qHSC, quiescent HSC; aHSC, activated HSC; cHSC, cycling HSC; iHSC, inflammatory HSC; cNeu, cycling neutrophil; Neu, neutrophil; Mono, monocyte; Baso, basophil, Mast, mast cell; Metab., metabolic intermediate. Results are from isolated MPP3 cultured for 6 hours with or without (±) LPS and Pam3CSK4 (L/P) stimulation. (**B**) Slingshot trajectory analysis of un-stimulated/stimulated MPP3 scRNAseq dataset with identified branches. (**C**) Gene ontology (GO) analysis of aHSC/iHSC-like (c8) cluster genes. (**D**) Separate UMAP projection of un- stimulated and stimulated MPP3 scRNAseq dataset with preferential trajectories. (**E**) GO analysis of qHSC-like (c0) cluster genes up-regulated upon stimulation (> 4-fold increase).

Previously defined marker genes (*8, 19*) allowed the identification of 5 immature (c0, c1, c2, c5, c9), 2 inflammatory (c7, c8), 2 intermediate (c3, c11) and 4 myeloid-primed (c4, c6, c10, c12) clusters (fig. S4A). Among the last, c4 and c6 were neutrophil-primed, c10 neutrophil/monocyte-primed, and c12 mast cell/basophil-primed clusters, while neither lymphoid, megakaryocyte, or erythroid lineage identity was detected in any clusters consistent with the myeloid-biased properties of MPP3. Slingshot (*26*) trajectory analyses identified 3 differentiation branches all starting from quiescent HSC (qHSC)-like c0, with branch 1 representing activation towards a cycling MPP state, branch 2 commitment towards myeloid differentiation via metabolically activated intermediates, and branch 3 the emergence of inflammatory subsets following stimulation (Fig. 2B). Gene ontology (GO) (*27*) analyses demonstrated enrichment in immune response genes in iHSC-like c8, metabolic process genes in metabolic intermediate c3, and myeloid differentiation genes in myeloid-primed c6, with cell cycle distribution further confirming the progressive activation status of these various MPP3 subsets (Fig. 2C, fig. S4, B to D, and table S2). Short-term 6 hours inflammatory stimulation particularly amplified the inflammatory branch 3 (c9, c8, c7) commitment, with up-regulation of inflammatory response genes already observed in stimulated qHSC-like c0 (Fig. 2, D and E). Altogether, these results deconstruct the molecular heterogeneity of the MPP3 compartment with constitutive cell cycle activation and myeloid differentiation trajectories, and inducible production of inflammatory subsets in response to stimulation.

### Secretory MPP3 stimulate myelopoiesis

We next investigated the function and regulation of this secretory ER^high^ MPP3 subset. Strikingly, ER^high^ MPP3 were almost exclusively Fc*γ*R^+^ but still differed from GMPs in their expression of other HSPC markers (Fig. 3A and fig. S5A). Conversely, the Fc*γ*R^+^ fraction of MPP3, which corresponds to the myeloid-primed clusters in scRNAseq analyses, was almost entirely ER^high^ (Fig. 3A and fig. S5B). In fact, a further harmonized scRNAseq UMAP directly projected ER^high^ MPP3 into *Fcgr3*-expressing myeloid-primed clusters and ER^low^ MPP3 into immature-like clusters (Fig. 3B and fig. S5C). Bulk RNA-sequencing and principal component (PC) analyses confirmed the similarity of ER^low^ MPP3 with HSCs and ER^high^ MPP3 with GMPs, with unfractionated MPP3 having a mixed gene identity (Fig. 3C). K-means clustering analyses further showed ER^high^ MPP3 variable genes clustered together with GMP variable genes and ER^low^ MPP3 variable genes clustered with HSC variable genes, with high expression of mature neutrophil genes such as *Mpo* and *Elane* (*18*) in ER^high^ MPP3 and immature HSC genes like *Gata2* and *Mpl* (*28, 29*) in ER^low^ MPP3 (fig. S5, D and E, and table S3). To probe the myeloid differentiation potential of MPP3 subsets, we next performed a series of *in vitro* and *in vivo* functional investigations. Compared to ER^low^ MPP3, ER^high^ MPP3 had significantly lower colony-forming capacity in both methylcellulose and single cell differentiation assays in liquid culture, which was also more restricted to myeloid differentiation, as well as faster division kinetics akin to GMPs (Fig. 3D and fig. S6, A and B). UMAP harmonization of our scRNAseq data with previously published myeloid progenitor profiling (*7*) directly projected ER^high^ MPP3 together with GMPs, while ER^low^ MPP3 appeared more related but still distinct from the transitional CMP population and lineage-committed MEP population (fig. S6C). Short-term *in vivo* lineage tracing in sub-lethally irradiated recipients directly confirmed that the entire reconstitution potential of MPP3 was provided by ER^low^ MPP3, with ER^high^ MPP3 barely registering for peripheral blood production (Fig. 3E). In fact, ER^low^ MPP3 (identified as the Fc*γ*R^-^ MPP3) were capable of producing ER^high^ MPP3 (identified as Fc*γ*R^+^ MPP3) both *in vitro* in short-term differentiation assays in liquid culture and *in vivo* following short-term infusion in recipient mice (fig. S6, D and E). Fc*γ*R^-^/ER^low^ MPP3 also significantly contributed to GMP production *in vivo*, likely through production of quickly differentiating Fc*γ*R^+^/ER^high^ MPP3 as well as by directly giving rise to CMPs (fig. S7A). To test the role of MPP3 subsets in myeloid regeneration *in vivo*, we then used an established anti-Ly6G depletion model that we previously showed induced transient MPP3 expansion prior to GMP and myeloid cell expansion (*12*). Remarkably, we found that the expansion of regenerative MPP3 resulted exclusively from an increase in Fc*γ*R^+^/ER^high^ MPP3, highlighting the role of this secretory subset as an amplification compartment in myelopoiesis (fig. S7B). Taken together, these results demonstrate that myeloid-primed Fc*γ*R^+^/ER^high^ MPP3 are a short-lived transitional population toward GMP commitment, while immature Fc*γ*R^-^/ER^low^ MPP3 represent the true multipotent part of the MPP3 compartment also capable of generating CMP and other lineage fates.

**Fig. 3.**
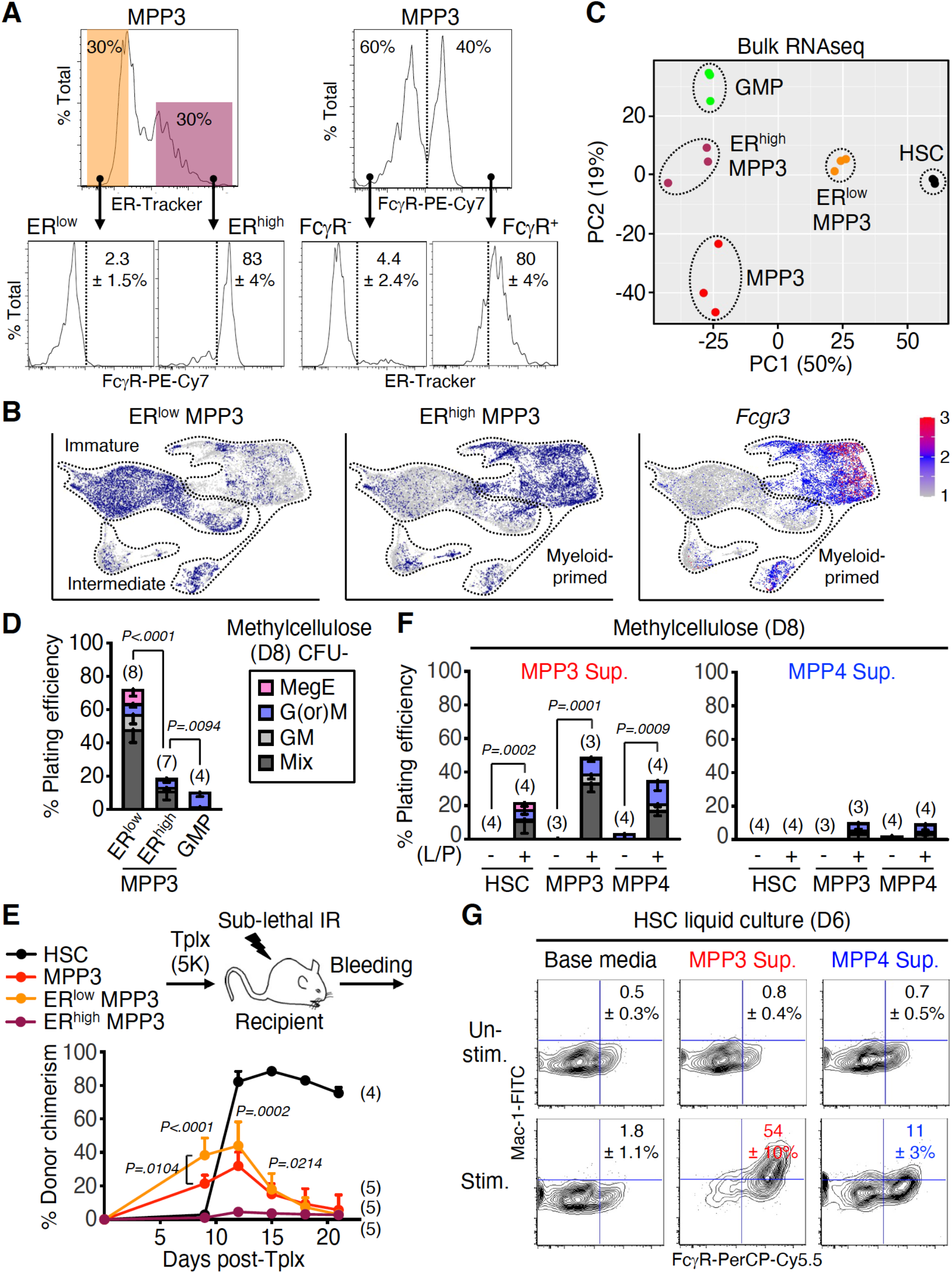
FcγR^+^/ER^high^ MPP3 subset is a transitional population regulating myelopoiesis. (**A**) Characterization of MPP3 subsets with representative FACS plots and quantification (n = 3) of Fc*γ*R^+^ frequency in ER^low^/ER^high^ MPP3 (left) and ER^high^ frequency in Fc*γ*R^-^/Fc*γ*R^+^ MPP3 (right). (**B**) Projection of ER^low^ MPP3 (left), ER^high^ MPP3 (middle) and *Fcgr3* expression (right) on the UMAP of harmonized un-stimulated/stimulated ER^low^/ER^high^/total MPP3 scRNAseq datasets. (**C**) Principal component (PC) analysis of HSC, MPP3, ER^low^ MPP3, ER^high^ MPP3 and GMP bulk RNAseq dataset. (**D**) Myeloid differentiation of ER^low^ MPP3, ER^high^ MPP3 and GMP in methylcellulose assays scored after 8 days (D); Mix, mixture of all lineages; GM, granulocyte/macrophage; G(or)M, granulocyte or macrophage; MegE, megakaryocyte/erythrocyte; CFU, colony-forming unit. (**E**) Short-term *in vivo* lineage tracing assay with experimental scheme for the transplantation (tplx) of 5,000 cells into each sub-lethally-irradiated (IR) recipient, and quantification of donor chimerism in peripheral blood over time. Significance was calculated between mice transplanted with ER^low^ or ER^high^ MPP3 unless otherwise indicated. (**F** and **G**) Myeloid differentiation effect of MPP3 and MPP4 supernatants (Sup.) on naïve HSCs, MPP3 and MPP4 plated either in (F) methylcellulose containing 20% supernatant or (G) in liquid cultures in 100% supernatant. Supernatants were collected upon culture of 10,000 MPP3 or MPP4 for 24 hours in 150 µl base media with or without (±) LPS/Pam3CSK4 (L/P) stimulation. Methylcellulose assays were scored after 8 days (D8), and differentiating cells in liquid cultures were analyzed by flow cytometry after 6 days (D6). Representative FACS plots and quantification (n = 3) of Mac- 1^+^/Fc*γ*R^+^ myeloid cell frequencies are shown for (G); Un-stim., un-stimulated; Stim., stimulated. Data are means ± S.D., and significance was assessed by a two-tailed unpaired Student’s t-test.

To understand the role of Fc*γ*R^+^/ER^high^ MPP3 secreted cytokines, we used supernatants from MPP3 and MPP4 cultured for 24 hours *in vitro* with or without LPS/Pam3CSK4 stimulation to initiate differentiation assays with naïve cells in methylcellulose and liquid culture (fig. S7C). Remarkably, supernatants from stimulated MPP3 induced myeloid colony formation in the range of full cytokine stimulation not only from naïve HSCs but also from naïve MPP3 and MPP4 (Fig. 3F and fig. S7C). In contrast, supernatants from stimulated MPP4 barely elicited myeloid differentiation from any of the naïve populations. Myeloid lineage tracing experiments provided similar results with only supernatants from stimulated MPP3 inducing myeloid differentiation from naïve HSCs with robust production of Mac-1^+^/Fc*γ*R^+^ myeloid cells (Fig. 3G). Collectively, these results identify a novel self-reinforcing regulatory function for a subset of secretory MPP3 that controls myelopoiesis through lineage-priming toward GMP differentiation and cytokine production to enhance myeloid cell production from other HSPC populations (fig. S7D).

### Expansion of secretory MPP3 in myeloid leukemia

Finally, we investigated the role of this newly identified Fc*γ*R^+^/ER^high^ MPP3 subset in malignant myelopoiesis. We used our inducible *Scl-tTA:TRE-BCR/ABL* (*BA^tTA^*) mouse model of myeloproliferative neoplasm (MPN), which we previously characterized for HSPC remodeling and MPP3 expansion associated with leukemic myeloid cell production (*10–12*). Strikingly, we found that the leukemic MPP3 expansion in *BA^tTA^* mice was entirely driven by an increase in Fc*γ*R^+^/ER^high^ MPP3 (Fig. 4, A and B). To gain molecular insights, we performed scRNAseq analyses on MPP3 isolated from 11-13 week-old aged-matched control (Ctrl) and diseased *BA^tTA^* mice (fig. S8A). Harmonized UMAP and Slingshot trajectory analyses identified 2 differentiation paths starting in immature HSC-like (c1) and progressing through cell cycle activation, with branch 1 representing myeloid differentiation, and branch 2 metabolic activation, the latter being specific to leukemic *BA^tTA^* MPP3 and positioned as an intermediate in the continuum from immature to *Fcgr3*-expressing myeloid-primed subsets (Fig. 4, C and D, and fig. S8, B and C). The leukemic-specific (c3, c8) cluster exhibited hallmark features of increased metabolism and biosynthesis processes, which were distinct from metabolic activation of normal MPP3 in culture and upon inflammatory stimulation (fig. S8, D to F). Together, these molecular data demonstrate the amplification of a unique subset of metabolically activated Fc*γ*R^+^/ER^high^ MPP3 in leukemic conditions, likely as a direct consequence of BCR/ABL activity (*30–32*).

**Fig. 4.**
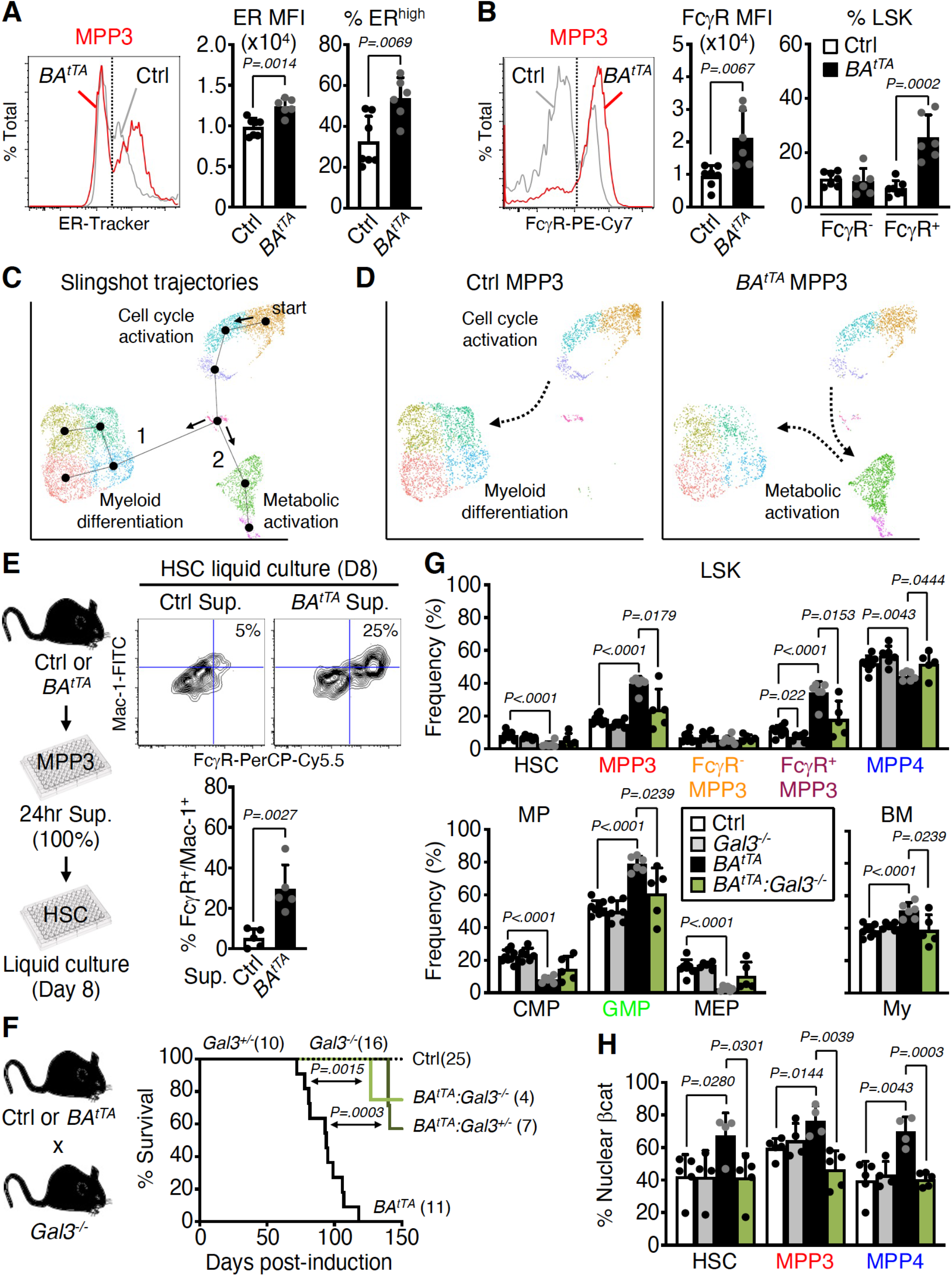
Constant secretion and myeloid amplification from leukemic FcγR^+^/ER^high^ MPP3 subset. (**A** and **B**) Quantification of (A) ER content and (B) Fc*γ*R surface expression in control (Ctrl) and *BA^tTA^* MPP3; MFI, mean fluorescence intensity. (**C**) Slingshot trajectory analysis of Ctrl/*BA^tTA^* MPP3 scRNAseq dataset with identified branches. (**D**) Separate UMAP projection of Ctrl and *BA^tTA^* MPP3 scRNAseq dataset with preferential trajectories. (**E**) Pro-myeloid differentiation effect of *BA^tTA^* MPP3 supernatant (Sup.) on naïve HSCs analyzed after 8 days (D) of liquid culture. Experimental scheme, representative FACS plots and quantification of Mac- 1^+^/Fc*γ*R^+^ frequencies are shown. Supernatants were collected upon culture of 10,000 Ctrl or *BA^tTA^* MPP3 for 24 hours in 150 µl base media. (**F**) Survival curve of *BA^tTA^* mice with Galectin 3 (*Gal3*) deletion. Results from Ctrl, *Gal3^+/-^*, *Gal3^-/-^*, *BA^tTA^*, *BA^tTA^:Gal3^+/-^* and *BA^tTA^:Gal3^-/-^* mice are shown; induction, Dox withdrawal. Significance was assessed by a Mantel-Cox test. (**G**) Changes in population size for HSPCs, myeloid progenitors and mature myeloid cells (My, Mac-1^+^/Gr-1^+^) in 11-13 week-old age-matched Ctrl, *Gal3^-/-^*, *BA^tTA^* and *BA^tTA^:Gal3^-/-^* mice. Results are expressed as percent of Lin^-^/Sca-1^+^/c-Kit^+^ (LSK), Lin^-^/Sca-1^-^/c-Kit^+^ (MP), and BM cells and are from 5 independent cohorts. (**H**) Quantification of nuclear *β*-catenin (*β*cat) positive HSC, MPP3 and MPP4 in a subset of age-matched Ctrl, *Gal3^-/-^*, *BA^tTA^* and *BA^tTA^:Gal3^-/-^* mice shown in (G). Data are means ± S.D., and significance was assessed by a two-tailed unpaired Student’s t-test except when indicated.

Next, we investigated the secretory activity of leukemic MPP3 by treating Ctrl and *BA^tTA^* MPP3 isolated from 11-13 week-old age-matched mice with or without LPS/Pam3CSK4 stimulation, and collecting supernatants after 24 hours for secretome analyses. Both Ctrl and *BA^tTA^* MPP3 secreted similar levels of cytokines upon stimulation with largely overlapping profiles and pro-myeloid differentiation effect on naïve HSCs in liquid culture (fig. S9, A to C). In contrast, *BA^tTA^* MPP3 constitutively secreted a unique set of 6 cytokines, IL-6, MIP1*α*, Galectin 1 (Gal1), Galectin 3 (Gal3) Lipocalin 2 (LCN2), and Granzyme B (GrB) (fig. S9D and table S4), some with well-known functions in myelopoiesis (*33*). In fact, un-stimulated *BA^tTA^* MPP3 supernatant increased the proliferation of naïve HSC, MPP3 and MPP4 and enhanced myeloid differentiation from naïve HSCs in liquid culture, although to a lower extent and with longer kinetics than LPS/Pam3CSK4 stimulated supernatants (Fig. 4E and fig. S9E). Except for LCN2, the cytokines constitutively secreted by *BA^tTA^* MPP3 were also found at elevated levels in the BM fluid of *BA^tTA^* mice (fig. S9, F and G and table S5), suggesting that they could play active roles in disease pathogenesis. Among those, Gal3 has been found highly expressed in various cancers and is a negative prognostic factor for AML patients (*34–36*). To test the importance of Gal3 secretion for myeloid amplification by leukemic MPP3, we used previously published *Gal3^-/-^* mice (*37*) to cross with *BA^tTA^* mice. *Gal3^-/-^* mice already showed reduced commitment towards myelopoiesis at steady state, with decreased HSC, Fc*γ*R^+^ MPP3, and GMP compartments, which was further exacerbated upon anti-Ly6G depletion treatment with impaired Fc*γ*R^+^ MPP3 and GMP expansion in regenerative conditions (fig. S10, A and B). Remarkably, both heterozygote and homozygote *Gal3* deletion significantly extend the survival of *BA^tTA^* mice with reversion of the leukemic expansion of Fc*γ*R^+^ MPP3, GMP and mature myeloid cells, and restoration of the defective production of MPP4 in *BA^tTA^:Gal3^-/-^* mice (Fig. 4, F and G). Gal3 is known to activate Wnt/*β*-catenin signaling by inhibiting GSK3*β* (*38, 39*), and we previously identified high Wnt/*β*-catenin activity in HSPCs as one of the key mechanisms driving MPP3 expansion and increased myelopoiesis in regenerative and leukemic conditions (*12*). Consistently, we found a striking reduction in nuclear *β*-catenin levels in *BA^tTA^:Gal3^-/-^* HSCs likely due to increased GSK3*β*activity caused by the loss of Gal3- mediated regulation (Fig. 4H). In fact, treatment of *BA^tTA^:Gal3^-/-^* HSCs with a GSK3*β* inhibitor restored aberrantly high nuclear *β*-catenin levels in leukemic HSCs (fig. S10C). Altogether, these results demonstrate that constitutive cytokine secretion by leukemic MPP3 plays a key role in enhancing myeloid lineage trajectory and amplifying myeloid cell production, thereby contributing to disease progression. They also identify a pivotal role for Gal3 secreted by *BA^tTA^* MPP3 in increasing Wnt activity in leukemic HSPCs, which could be therapeutically targeted to dampen engagement of emergency myelopoiesis pathways in leukemic conditions.

## Discussion

Our results uncovered a previously unappreciated self-regulatory mechanism in the HSPC compartment mediated by a secretory subset of myeloid-biased multipotent progenitors (MPP3) that controls myeloid commitment through lineage priming and amplifies “on demand” myeloid cell production via autocrine and paracrine effects in the local BM niche microenvironment (fig. S10D). In response to an acute inflammatory challenge, the MPP3 compartment is transiently remodeled via the production of inflammatory intermediates to increase the numbers and trigger the secretion of pro-myeloid differentiation cytokines from Fc*γ*R^+^/ER^high^ MPP3. During MPN development, the MPP3 compartment is re-structured with amplification of metabolically-activated intermediates that fuel the overproduction of Fc*γ*R^+^/ER^high^ MPP3, which in turn constitutively secrete cytokines reinforcing both myeloid commitment and differentiation. It now remains to be determined whether other leukemia-initiating events will also amplify malignant myeloid cell production through changes in MPP3 lineage priming and cytokine secretion. Although with different modalities in stress and disease conditions, MPP3 act in both cases as a self-reinforcing compartment of myeloid amplification and a key regulator of myelopoiesis. This identify MPP3 as one of the first myelopoiesis-regulatory population (or MyReg) that act similarly to Tregs but for the myeloid lineage and to control stem and progenitor cell fate. In this context, it is likely that MPP3 will play an important role in amplifying myeloid cell production in other deregulated contexts such as infectious disease conditions and aging. Considering the autocrine and paracrine effects of MPP3-secreted cytokines in tailoring HSCs and other progenitor fate, it is also likely that MPP3 function will underlie some, if not most, of the cell-autonomous HSC behaviors in single cell transplantation experiments. Collectively, our findings identify a novel mechanism regulating myelopoiesis at the early stage of hematopoietic commitment, which represents an ideal cellular compartment to target therapeutically to rebalance lineage output in stress and disease conditions.

## Acknowledgements

We thank Dr. M. Ansel (UCSF) for *Blimp-1-Yfp* mice, Dr. J. Pereira (Yale) and IsoPlexis for help with confirmatory single cell secretion experiments, M. Kwak (Yale) for preparation of antibody barcode glass slides, Dr. A. Collins (CUIMC) for HSC bulk RNAseq data, J. Wong (Gladstone Institute at UCSF) for electron microscopy analyses, M. Lee (UCSF) and M. Kissner (CUIMC) for management of our Flow Cytometry Core facilities, M. Kanai (CUIMC) for technical assistance, and all members of the Passegué laboratory both at UCSF and CUIMC for critical insights and suggestions.

## Author contributions

Conceptualization: YK, EP

Methodology: YK, SZ, JC, EP

Investigation: YK, HP, SZ, JC, MW

Formal Analysis: YK, HP, SZ, JC

Visualization: YK, EP

Funding acquisition: YK, EP

Project administration: YK, EP

Supervision: RF, EP

Writing – original draft: YK, EP

Writing – review & editing: YK, EP

## Data and materials availability

Data sets that support the findings of this study have been deposited in the Gene Expression Omnibus (GSE181902). All data are available in the main text or the supplementary materials. Correspondence and requests for materials should be addressed to EP.

## Supplementary Materials

### Materials and Methods

#### Mice

All mice were bred and maintained in mouse facilities at UCSF or CUIMC in accordance with IACUC protocols approved at each institution. CD45.2 C57BL/6J (000664), CD45.1 C57BL/6- BoyJ (002014) and B6.Cg-*Lgals3^tm1Poi^*/J (006338) mice were purchased from the Jackson Laboratory. *Prdm1-Yfp* mice were obtained from Dr. Mark Ansel (UCSF)^40^. *β-actin-Gfp*, *Scl-tTA:TRE-BCR/ABL*, and *Tnfa^-/-^* mice were previously described^2,10,14^. *BA^tTA^:Gal3^-/-^* mice were obtained by breeding B6.Cg-*Lgals3^tm1Poi^*/J mice with *Scl-tTA:TRE-BCR/ABL* mice. Respective wild type (WT) littermates or single transgenic animals were used as controls (Ctrl). Six- to 12- week-old mice were used as donor for cell isolation, and 8- to 12-week-old congenic mice were used as recipients for transplantation experiments. For BCR/ABL induction, mice were withdrawn from doxycycline (Dox) containing water at 5 week of age. No specific randomization or blinding protocol was used, and both male and female animals were used indifferently in the study.

#### Flow Cytometry

Staining of hematopoietic cells were performed as described previously^12^. In brief, BM cells were obtained by crushing leg, arm, and pelvic bones in staining media composed of Hanks’ buffered saline solution (HBSS) containing 2% heat-inactivated FBS (Corning, 35-011-CV). Red blood cells (RBC) were removed by lysis with ACK (150 mM NH4Cl/10 mM KHCO3) buffer, and single-cell suspensions of BM cells were purified on a Ficoll gradient (Histopaque 1119, Sigma-Aldrich). Spleens were mechanically dissociated in staining media and ACK lysed to remove contaminating RBCs. Blood was collected in ACK buffer containing 10% EDTA from intra-orbital bleed and further lysed in ACK buffer to remove contaminating RBCs. Cellularity was determined by ViCELL-XR automated cell counter (Beckman-Coulter). For HSC and progenitor isolation, BM cells were pre-enriched for c-Kit^+^ cells using c-Kit microbeads (Miltenyi Biotec, 130-091-224) and an AutoMACS cell separator (Miltenyi Biotec). Unfractionated or c-Kit-enriched BM cells were then incubated with purified rat anti-mouse lineage antibodies (CD3, BioLegend, 100202; CD4, eBioscience, 16-0041-82; CD5, BioLegend, 100602; CD8, BioLegend, 100702; CD11b, BioLegend, 101202; B220, BioLegend, 103202; Gr1, eBioscience, 14-5931-85; Ter119, BioLegend, 116202) followed by goat anti-rat-PE-Cy5 (Invitrogen, A10691) and subsequently blocked with purified rat IgG (Sigma-Aldrich). Cells were then stained with Sca-1-PB (BioLegend, 108120), c-Kit-APC-Cy7 (BioLegend, 105826), CD48-A647 (BioLegend, 103416), CD150-PE (BioLegend, 115904), and Flk2-Bio (eBioscience, 13-1351-85) followed by SA- BV605 (BioLegend, 405229), CD34-FITC (eBioscience, 11-0341-85) and FcγR-PE-Cy7 (BioLegend, 101318). For ER-Tracker staining, BM cells were stained with 0.3 μM ER-Tracker green (Invitrogen, E34251) in Ca^2+^/Mg^2+^-containing HBSS (Gibco, 14025092) for 15 min at 37°C in a 5% CO2 water jacket incubator. For plasma cell isolation, spleen cells from *Prdm1-Yfp* mice were stained with B220-APC-eFluor780 (eBioscience, 47-0452-82) and CD138-APC (BD Pharmingen, 558626). For *in vitro* myeloid differentiation of naïve HSCs, cultured cells were stained with Mac-1-FITC (eBioscience, 11-0112-82) and FcγR-PerCP-eFluor710 (eBioscience, 46-0161-82). For *in vitro* single cell myeloid lineage assays, cultured cells were stained with CD45-APC-Cy7 (BD Pharmingen, 557659), CD71-BUV395 (BD Pharmingen, 740223), CD41- BV510 (BioLegend, 133923), CD61-PE (Biolegend, 104308), Gr-1-BV421 (Biolegend, 108433), Mac-1-PE-Cy7 (eBioscience, 25-0112-82), FcεRI-APC (BioLegend, 134316). For donor-derived chimerism analyses in transplanted mice, blood cells were stained with Gr-1-eFluor450 (eBioscience, 48-5931-82), Mac-1-PE-Cy7, B220-APC-eFluor780, CD3-eFluor660 (eBioscience, 50-0032-82), Ter-119-PE-Cy5 (eBioscience, 15-5921-83), CD45.1-PE (eBioscience, 12-0453-83) and CD45.2-FITC (eBioscience, 11-0454-85). For short-term *in vitro* Fc*γ*R^-^ MPP3 and Fc*γ*R^+^ MPP3 culture, cells were stained with Sca-1-PB, c-Kit-APC-Cy7, CD48-A647, CD150-PE, Mac- 1-FITC and FcγR-PE-Cy7. For *in vivo* infusion of *β-actin-Gfp* cells, c-Kit-enriched recipient BM cells were stained with lineage cocktails as described above and then stained with Sca-1-PB, c-Kit-APC-Cy7, CD48-A647, CD150-PE, FcγR-PE-Cy7 and CD34-Bio (BioLegend, 119304) followed by SA-BV605 (BioLegend, 405229). Stained cells were finally re-suspended in staining media containing 1 µg/ml propidium iodide (PI) for dead cell exclusion. Cell isolations were performed on a Becton Dickinson (BD) FACS Aria II (UCSF) or FACS Aria II SORP (CUIMC) using double sorting for purity. Cell analyses were performed on a BD LSR II (UCSF), BD Celesta (CUIMC), Agilent Novocyte Quanteon (CUIMC) or Bio-Rad ZE5 (CUIMC) cell analyzer. All data were analyzed using FlowJo (Treestar).

#### In vitro assays

All cultures were performed at 37°C in a 5% CO2 water jacket incubator (Thermo Scientific) and, except for TNF*α*, all cytokines were purchased from PeproTech. Mouse TNFα was obtained from Genentech under an MTA. To harvest supernatants, cells (10,000 per well of a 96-well plate) were grown in 150 µl base media consisting of Iscove’s modified Dulbecco’s media (IMDM) (Gibco) with 5% FBS (StemCell Technology), 50 U/ml penicillin, 50 μg/ml streptomycin, 2 mM L-glutamine, 0.1 mM non-essential amino acids, 1 mM sodium pyruvate and 50 μM 2- mercaptoethanol, and containing only SCF (25 ng/ml), TPO (25 ng/ml) and Flt3-L (25 ng/ml) as cytokines. For stimulation, cells were cultured in the presence of 100 ng/ml LPS (Sigma-Aldrich, L4391) and 1 μg/ml Pam3CSK4 (R&D Systems, 4633) for up to 24 hours. For testing supernatant effects in liquid culture, naïve HSCs (1,000 cells per well of a 96-well plate) were grown in 150 µl of 100% harvested supernatants for either 6 days without any media replacement or 8 days with replenishing half of the culture supernatants every 3 days. For CFSE dilution assay, naïve cells (2,000 cells per well of a 96-well plate) were labeled with 2.5 μM CFSE (Molecular Probes, C1157) as described previously^12^ and cultured for 3 days in 150 µl of 100% supernatant. For other liquid culture assays, cells (2,000 to 3,000 per well of a 96 well plate) were grown in 200 µl base media supplemented with IL-11 (25 ng/ml), IL-3 (10 ng/ml), GM-CSF (10 ng/ml) and EPO (4 U/ml) for full cytokine cocktail, media and cultured for indicated time before analyses. Depending on experiments, 2 μM BMS-345541 (Sigma-Aldrich, B9935), 2 μM KN-93 (Sigma-Aldrich, K1385) and 30 µM CHIR 99021 (Selleckchem, S1263) was also added. To harvest supernatants for TNF*α* addition experiments, cells were exposed to 1 μg/ml TNF*α* (Genentech) in full cytokine cocktail to ensure proper cell viability. For single cell division counts, individual cells were directly sorted per well of a Terazaki plate containing 10 μl of full cytokine media, and the number of cells per well was manually counted under a microscope every 12 hours. For single cell differentiation assays, individual cells were directly sorted per well of a 96 well-plate in 200 µl of IMDM containing 10% FBS (StemCell Technology), 20% BIT 9500 (StemCell Technology, 9500), 5% PFHM II (Gibco, 12040077), 50 U/ml penicillin, 50 μg/ml streptomycin, 2 mM L-glutamine, 0.1 mM non-essential amino acids, 1 mM sodium pyruvate, 55 μM 2-mercaptoethanol, SCF (50 ng/ml), TPO (25 ng/ml), Flt3-L (10 ng/ml), IL-6 (10 ng/ml), IL-3 (10 ng/ml), GM-CSF (5 ng/ml) and EPO (4 U/ml). Small megakaryocytic colonies were manually counted under a microscope at day 5, and all other colonies were harvested and scored by flow cytometry analyses after 6 days of culture. For methylcellulose colony assays, cells were plated into 35-mm dish (100 cells/dish) containing 1 ml methylcellulose (StemCell Technologies, M3231) supplemented with 50 U/ml penicillin, 50 μg/ml streptomycin, and either the full cytokine cocktail described above or just 20% culture supernatants. Colonies were manually scored under a microscope after 8 days of culture.

#### In vivo assays

For granulocyte depletion, mice were injected once intraperitoneally with 0.1 mg of IgG control (clone 2A3) or anti-Ly6G antibody (BioXcell, BP0075-1) in 200 µl PBS. For short-term *in vivo* lineage tracing assays, CD45.1 recipient mice were sub-lethally irradiated (8.5 Gy, delivered in split doses 3 hours apart) using an X-ray irradiator (Faxitron) and injected retro-orbitally with 5,000 CD45.2 donor cells within the next 6 hours. Irradiated recipient mice were administered polymyxin/neomycin-containing water for 4 weeks following transplantation to prevent opportunistic infection, and analyzed over time by repeated bleedings. Peripheral blood (PB) was obtained from retro-orbital plexus, and collected in tubes containing 4 ml of 10 mM EDTA/ACK lysis buffer for flow cytometry analyses. For short-term *in vivo* differentiation assays, 10,000 donor cells isolated from *β-actin-Gfp* mice were retro-orbitally infused into recipient mice, which were analyzed for BM contribution at the indicated times.

#### Cytokine analyses

Culture supernatants were clarified by filtering through 0.22 μm filter (Millipore, SLGV004SL) and stored at -80°C until use. BM fluids were prepared as previously decribed^12^ and stored at - 80°C until use. For ELISA measurements, 50 µl of culture supernatants were analyzed with IL-6 (eBioscience, 50-172-18) or TNF*α* (eBioscience, 88-7324-22) ELISA kits according to the manufacturer’s instructions. For Luminex cytokine multiplex bead arrays, 25 µl of culture supernatants were analyzed for a custom-made panel of 5 cytokines (IL-1*β*, IL-6, GM-CSF, TNF*α*, MIP1*α*; Invitrogen, PPX-05) according to the manufacturer’s instructions. For Raybiotech 200 mouse cytokine arrays, 500 µl of pooled culture supernatants or BM fluids were sent per sample to Raybiotech for quantitative proteomics services using Mouse Cytokine Array Q4000 kit. Quantile normalization was performed for direct comparison of secreted cytokine profiles from different array experiments using R Bioconductor package, and hierarchical clustering was conducted using pheatmap package of R (https://cran.r-project.org/web/packages/pheatmap/index.html). The statistical significance of the different distributions of secreted cytokine under diverse biological conditions was determined using Kruskal-Willis test (P-value < 0.05, R version 3.3: www.r-project.org).

#### Single cell secretion assays

Polydimethylsiloxane (PDMS) microwell array and antibody barcode glass slides were prepared and screened with FITC-labelled BSA as described^41^, then blocked with 3% BSA for 1 hour at room temperature (RT) and rinsed with base media just before usage. The PDMS microwell array was placed facing upward and the media removed until just a thin layer remained on the array surface. Cell suspension (25,000 cells in 100 µl base media ± LPS/ Pam3CSK4 stimulation) was pipetted onto the microwell array and allowed to settle for 10 min. The antibody glass slide was then put on top of the PDMS microwell array, with antibody barcode resting on the cell capture chambers, clamped tight together and imaged on an automatic microscope stage to acquire optical images recording the number and location of trapped cells in each microwell. Following imaging, the assembly was incubated for 18 hours at 37°C in a 5% CO2 water jacket incubator, after which the antibody barcode glass slide was removed and submitted to ELISA immunoassay detection procedure. In brief, a mixture of biotinylated detection antibodies was pipetted onto the glass slide and incubated for 45 min at RT followed by washing with 3% BSA solution. APC dye-labeled streptavidin (eBioscience, 17-4317-82) was added onto the glass slide and incubated for 30 min at RT, then washed with 3% BSA again and blocked with 3% BSA for 30 min at RT. The glass slide was then dipped sequentially in 2 Dulbecco’s PBS baths and 2 DI water baths before being finally blown dry. Genepix 4000B (UCSF) and 4200A (Yale) scanners (Molecular Devices) were used to obtain scanned fluorescent images for FITC (488 nm) and APC (635 nm) channels. Immunofluorescence and optical images were analyzed with GenePix Pro software (Molecular Devices) to align the microwells array template and extract fluorescence intensity values for wells determined to contain only a single cell. Fluorescence data were used to generate heatmap and scatterplots with Excel (Microsoft) and Prism (GraphPad).

#### Immunofluorescence staining of cells

Isolated cells (2,000-3,000 in 5-10 µl of IMDM) were pipetted onto poly-L-lysine coated slides (Sigma-Aldrich, P0425-72EA), settled down for 15 min at RT, fixed with 4% PFA for 10 min at RT, then washed 3 times with PBS, and permeabilized/blocked for 1 hour at RT with 0.1% Tween- 20 in 10% FBS (Corning) in PBS, which was then used as antibody incubation buffer for all the subsequent steps. Cells were incubated overnight at 4 °C with a mouse monoclonal anti-KDEL (Abcam, ab12223) or a rabbit anti-mouse β-catenin (Cell Signaling, 9582S) primary antibody, washed 3 times with PBS, and incubated for 1 hour at RT with a goat anti-mouse IgG A488 (Invitrogen, A11029) or a donkey anti-rabbit-A555 (Invitrogen, A31572) secondary antibody. Cells were then washed 3 times with PBS, stained with 1 μg/ml DAPI (Sigma-Aldrich, 32670) for 10 min at RT, washed 3 times with PBS and finally slides were mounted with VectaShield (Vector Laboratories, H-1000). Cells were imaged on a Nikon Ti Eclipse inverted confocal microscope for KDEL images (60x objective) and images were processed using Fiji (https://fiji.sc). Cells were imaged on an Olympus epifluorescence microscope (60x objective) for manual scoring of nuclear β-catenin staining. At least 100 cells per condition were randomly captured for quantification.

#### Immunofluorescence staining of bone sections

Femurs were embedded in OCT (Tissues-Tek, 4583), snap-frozen in a 100% ethanol/dry ice slurry, and kept at -80 °C until sectioning. Thin 7 μm section were obtained upon cryosection at -30°C with a CryoJane tape transfer system (Leica, 39475205) and a tungsten blade, and were kept at - 80°C before staining. Sections were first fixed with 100% acetone kept at -20°C for 10 min, dried for 5 min at RT, blocked for 90 min with 10% goat-serum (Gibco) in PBS and washes 3 times with PBS for 5 min at RT. The same wash procedure was also used in between each staining step performed in 10% goat-serum in PBS. For HSPC staining, sections were incubated with A488- conjugated CD48 (BioLegend, 103414) for 90 min at RT, blocked with 20 μg/ml Rat IgG (Sigma-Aldrich, I8015-10MG) for 10 min at RT, and then stained with PE-conjugated ESAM (BioLegend, 136203) and Alexa647-conjugated lineage markers CD3e (BioLegend, 100209), B220 (BioLegend, 136203), Gr-1 (Biolegend, 108418), Mac-1 (Biolegend, 101218), and CD150 (Biolegend, 115918) for 90 min at RT. Sections were then counterstained with 1 μg/ml DAPI in PBS for 10 min at RT and mounted with Fluoromount G (Southern Biotech, 0100-01) and imaged on an SP5 upright confocal microscopes (Leica) with 20x objectives. Images were processed and analyzed using Volocity software (Perkin Elmer v.6.2), Imaris (Bitplane v.8.2), and Photoshop (Adobe v.CS5). For MPP3 staining with the immunofluorescence scheme for *in situ* imaging, cells were stained by flow cytometry as described above and 1,000 MPP3 were directly deposited onto poly-L-lysine coated slides and fixed with 100% acetone for 5 min at -20°C. Of note, acetone fixation strip the flow cytometry surface markers allowing re-staining of isolated MPP3. Slides were then blocked for 90 min at RT with 10% goat serum in PBS, stained as described above for HSPC staining for tissue sections, and then mounted with VectaShield (Vector Laboratories, H- 1200) containing 1 μg/ml DAPI and imaged on SP5 upright confocal microscope with oil immersion 63x objective. Images were processed using Volocity software, and 100 cells were scored to construct a library of representative images.

#### Electron Microscopy

Cells (50,000 to 100,000 per sample) were fixed in 0.1M NaCacodylate (pH 7.4) containing 1% paraformaldehyde and 2% glutaraldehyde on ice for 30 min, and pelleted at 3000 x g for 10 min at 4°C. Samples were then submitted to the Gladstone Institute (UCSF) Electron Microscopy Core Facility for standard transmission electron microscopy (TEM) ultrastructural analyses. MPP3 images were manually scored for the presence or absence of cytoplasmic rough endoplasmic reticulum (ER) structures.

#### SAbiosciences UPR PCR array

Cells (5,000 per sample) were directly sorted into 500 μl Trizol LS (Life Technologies, 10296-010) and RNA was extracted using Arcturus PicoPure RNA Isolation kit (Applied Biosystems, KIT0204) according to the manufacturer’s instructions. RNA concentration was measured using a bioanalyzer and 1 ng of RNA was treated with DNase I (Invitrogen, 18068-015) and reverse-transcribed using SuperScript III kit and random hexamers (Invitrogen, 18080-051) to make cDNA. All cDNA samples were pre-amplified for pathway specific genes and PCR arrays were performed according to the manufacturer’s instructions (Qiagen, PAMM-089Z). 7900HT Fast Real-Time PCR system was used to run the array and the data were analyzed using the web-based SABiosciences RT² Profiler™ PCR Array Data Analysis software. Values were normalized to *Gusb* expression.

#### Bulk RNA sequencing

Cells (5,000 to 7,000 per sample) were directly sorted into 350 μl RLT lysis buffer containing 1% β-mercaptoethanol from the RNeasy Plus Micro Kit (Qiagen, 74034) and RNA was isolated according to the manufacturer’s instructions. RNA samples were submitted to the NYU Genome Technology Center for low input RNA sequencing. Briefly, RNA samples with RNA integrity number (RIN) > 9.5 as measured with a Bioanalyzer (Agilent Technologies) were subjected to further processing. Libraries were prepared using Trio RNA-seq Library Preparation Kit (NuGen, 0507) and sequenced on an Illumina NovaSeq 6000 for an average of ∼82.5 million paired reads per sample. The initial quality control of raw reads was performed using FastQC (https://www.bioinformatics.babraham.ac.uk/projects/ fastqc/) for read quality assessment, and HISAT2^42^, the spliced junction mapper, to assess read duplication and read quality. The reference gene model of HISAT2 was based on the Mus musculus genome (GRCm38). The abundance of transcript reads were estimated using Salmon^43^ and genes with low read counts were filtered out (<10 read counts across all samples). Differential gene expression analysis was carried out using R/Bioconductor DEseq2^44^ package. Normalized read counts were computed by dividing the raw read counts by size factors and fit to a negative binomial distribution. P-values were first corrected by applying empirical estimation of the null distribution using the R package and then adjusted for multiple testing with the Benjamini–Hochberg correction. Genes with an adjusted P-value less than 0.05 and fold change values greater than 2 were considered differentially expressed. Volcano plots were generated using the R (v.3.6.0) packages (ggplot2 and EnhancedVolcano). The enrichment of gene signatures based on KEGG pathway and GO term was examined using R package clusterProfiler^45^, and IPA tool (Ingenuity Pathway Analysis, Qiagen). For highly variable gene (HVG) selection, the background distribution of the coefficient of variation of gene expression was calculated using random permutation of gene expressions across samples. HVGs were then selected based on the calculated random distribution of the variance of gene expression (P-value < 0.05). Out of 19,639 genes, 981 genes were determined as HVGs of compared samples. With selected HVGs, K-means clustering approach identified the sequential pattern of gene expression changes across samples.

#### Droplet-based single cell RNA sequencing

For cultured samples, unfractionated MPP3 (50,000 cells per sample) and ER^high^/ER^low^ MPP3 subsets (30,000 cells per sample) were sorted per well of a 96 well-plate into 150 µl base media ± LPS/Pam3CSK4 stimulation and cultured for 6 hours. Cells were then washed and resuspended in HBSS/2% FBS at a concentration of 1000 cells/μl. For freshly isolated samples, MPP3 (30,000 to 50,000 cells per sample) were directly sorted in HBSS/2% FBS at a concentration of 1000 cells/μl. Samples were then submitted to the Columbia Genome Center Single Cell Analysis Core for microfluidic cell processing, library preparation and sequencing. Briefly, cell viability and concentration were measured using a Countess II FL Automated Cell Counter (Thermo). RNA-seq library was prepared using a Chromium Single Cell 3′ Library & Gel Bead Kit v2 (10X Genomics) according to the manufacturer’s instructions. Samples (15,000 cells per sample per condition) were loaded and sequenced on an Illumina HiSeq4000 sequencer for an average of ∼326 million reads per sample. Sequencing data were aligned and quantified using the Cell Ranger Single-Cell Software Suite (version 2.0, 10xGenomics) against the Mus musculus genome (GRCm38) provided by Cell Ranger. Using Seurat package of R (version 3)^25^, cells with fewer than 200 detected genes and of which the total mitochondrial gene expression exceeded 20% were removed. All of raw read data were passed the threshold of the proportion of doublet cells based on the DoubletFinder (<10% of doublet cells)^46^ and the number of gene expressions per cell (<7500 expressions per cell). Bioinformatic analysis was conducted based on diverse R package and python-based analysis. The clustering of the preprocessed scRNA-seq data was based on the UMAP (Uniform Manifold Approximation and Projection for Dimension Reduction) approach of the Seurat version3 package of R. The ‘FindAllMarker’ function of Seurat package was used to determine the list of cluster-defining genes, the ‘Dotplot’ function to determine the cellular identity of each cluster based on the expression profile of previously published ID genes^8,19^, the ‘CellCycleScoring’ function to infer cell cycle status of each cell based on regression approach, and the ‘FeaturePlot’ function to highlight a group of cells of interest on the UMAP representations. For comprehensive integration and comparison of scRNAseq data across samples from diverse experimental conditions, nearest neighbor integration was used. We inferred the trajectory of cell state transitions using slingshot^26^ of R package based on the analysis results of Seurat version3.

#### Quantitation and Statistical Analysis

All experiments were repeated as indicated; n indicates the numbers of independent biological repeats. Data are expressed as mean ± standard deviation (S.D.). Mice for treatment and transplantation were randomized, samples were alternated whenever possible, and no blinding protocol was used. Statistical significance was evaluated by a two-tailed unpaired Student’s t-test unless otherwise indicated. *P* values < 0.05 were considered statistically significant. Figures were made with GraphPad Prism software.

---

**Fig. S1.**
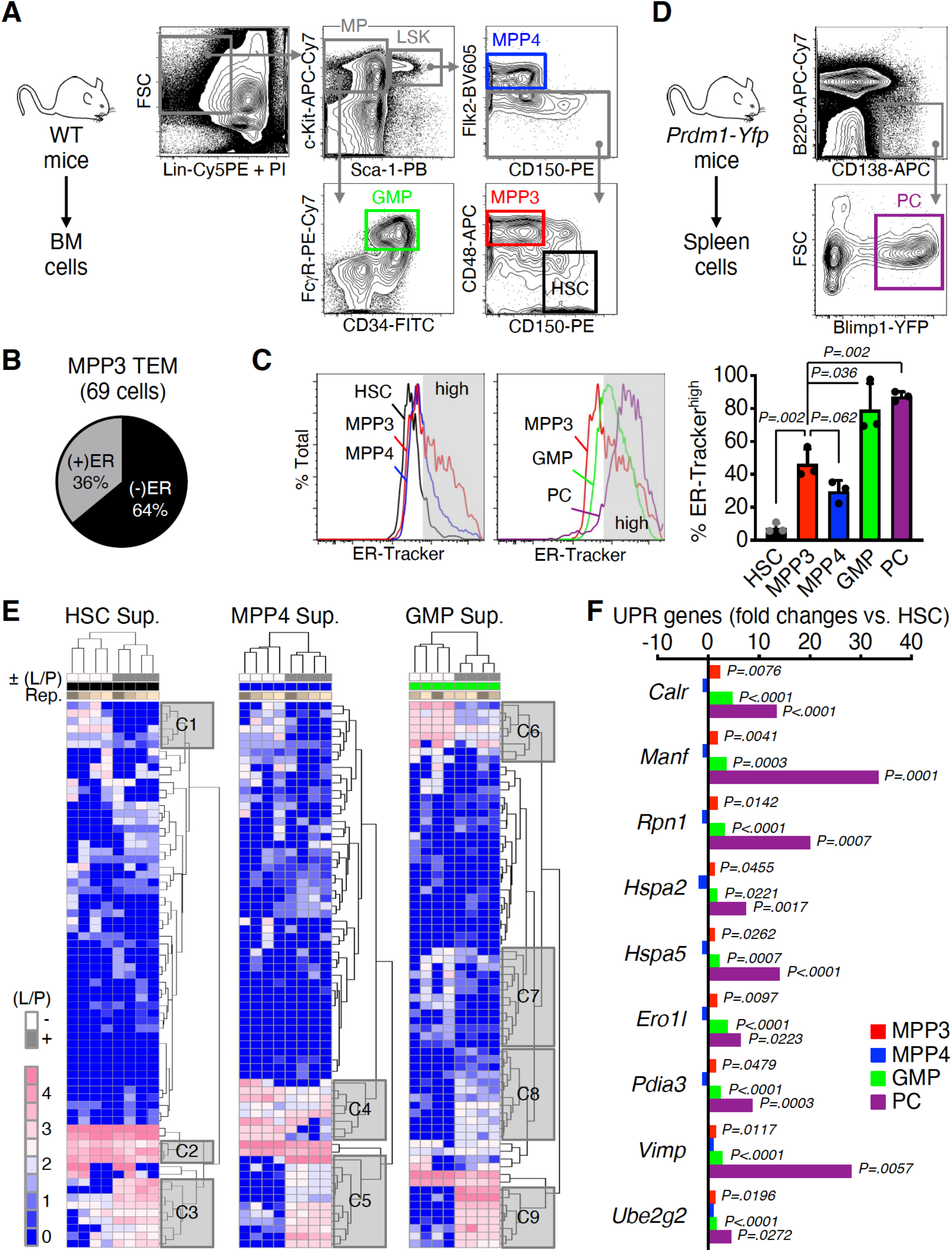
Secretory activity of HSPCs. (**A**) Gating strategy used for identifying and isolating bone marrow (BM) hematopoietic stem cells (HSC), multipotent progenitors (MPP3 and MPP4), and granulocyte/macrophage progenitors (GMP) from wild type (WT) donor mice. (**B**) Quantification of transmission electron microscopy (TEM) image of MPP3 with high (+) and low (-) endoplasmic reticulum (ER) volume. (**C**) ER-Tracker staining of HSPCs, GMPs, and plasma cells (PC) with representative FACS plots and quantification of ER-Tracker^high^ fraction (grey shaded area on histograms). Data are shown as means ± S.D. (**D**) Gating strategy used for identifying and isolating splenic plasma cells from Prdm1-Yfp mice. (**E**) Heatmap of unsupervised clustering of cytokines secreted in HSC, MPP4 and GMP supernatants after quantile normalization. Supernatants were collected upon culture of 10,000 cells for 24 hours in 150 µl base media with or without (±) LPS/Pam3CSK4 (L/P) stimulation; Rep., independent repeats. Representative clusters (C1 to C9) of secreted cytokines changed upon stimulation are provided in Extended Data Table 1. (**F**) SAbiosciences PCR array of unfolded protein response (UPR) genes in HSPCs, GMP and plasma cells (n = 3). Results are expressed as log2 mean fold expression relative to HSCs (set to 0). Significance was assessed by a two-tailed unpaired Student’s t-test.

**Fig. S2.**
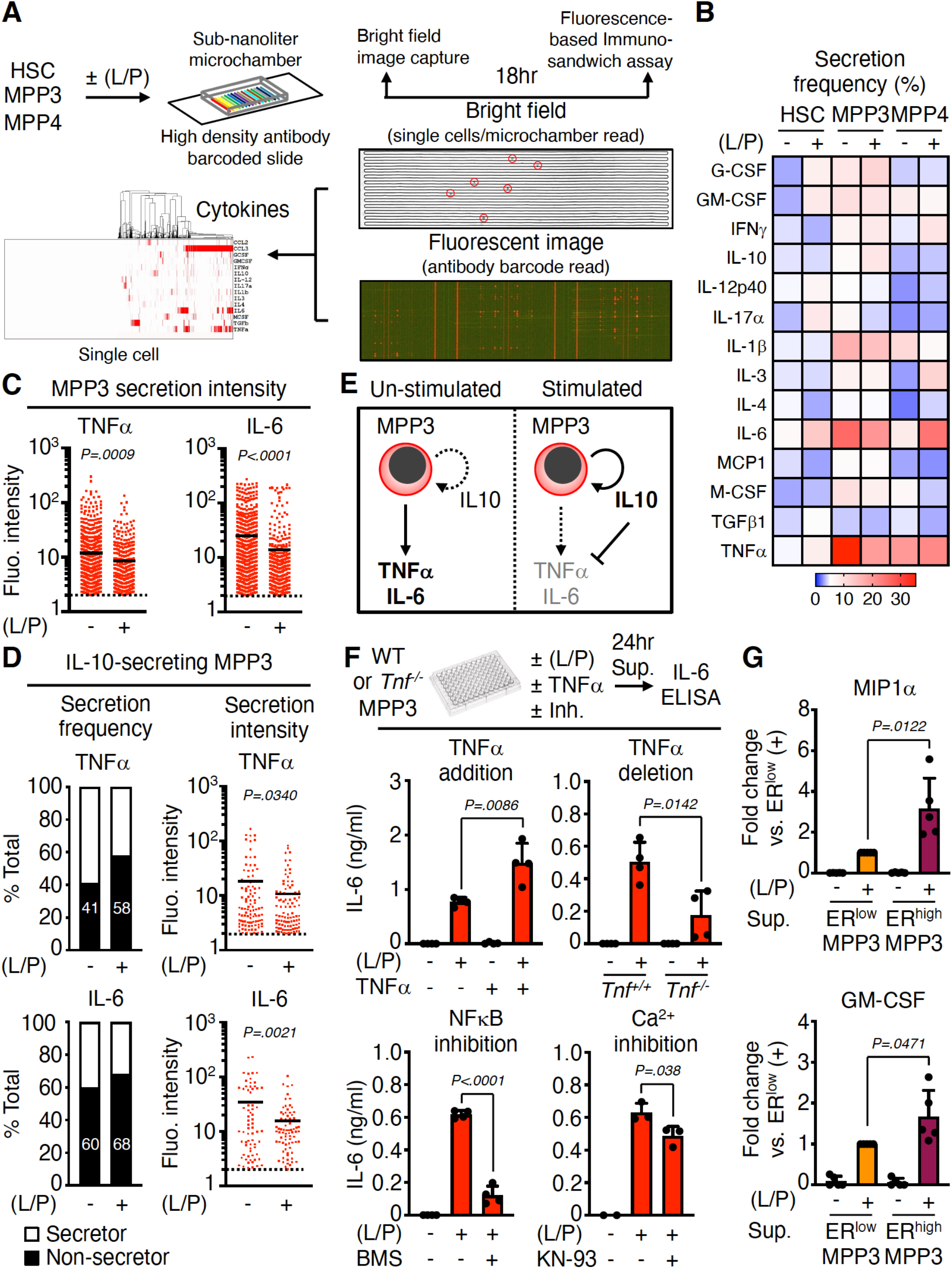
Autocrine effect of MPP3 secretion. (**A** to **E**) Secretory activity of HSPCs at the single cell level with: (A) experimental scheme of HSC, MPP3 and MPP4 single cell secretion assay with or without LPS/Pam3CSK4 (L/P) stimulation for 18 hours in culture (14 known cytokines were pre-selected for this assay); (B) heatmap of secretion frequency by single un-stimulated and stimulated HSC, MPP3, and MPP4; (C) TNF*α* and IL-6 secretion intensity by all secreting MPP3 with or without stimulation (cells with fluorescence (Fluo.) signal intensity above threshold (set to 2) are counted as secretors; black line, mean value); (D) TNF*α* and IL-6 secretion frequency and secreting intensity by IL-10-secreting MPP3; and (E) Model depicting the effect of IL-10 on TNF*α* and IL-6 secretion by individual MPP3. (**F**) Changes in IL-6 secretion by MPP3 upon TNF*α* addition (1 μg/ml), TNF*α* genetic deletion, and NF-*κ*B (BMS345541, 2 μM) or Ca^2+^ (KN- 93, 2 μM) signaling inhibition. Supernatants were collected upon culture of 10,000 MPP3 for 24 hours in 150 µl base media or full cytokine media (± TNF*α*) with or without (±) LPS/Pam3CSK4 (L/P) stimulation and the indicated inhibitor. Results are from ELISA measurements. (**G**) Differential secretion of MIP1*α* and GM-CSF by ER^high^ vs. ER^low^ MPP3 upon stimulation. Results are from 24 hours supernatants analyzed by Luminex cytokine bead array. Data are means ± S.D. except when indicated, and significance was assessed by a two-tailed unpaired Student’s t-test.

**Fig. S3.**
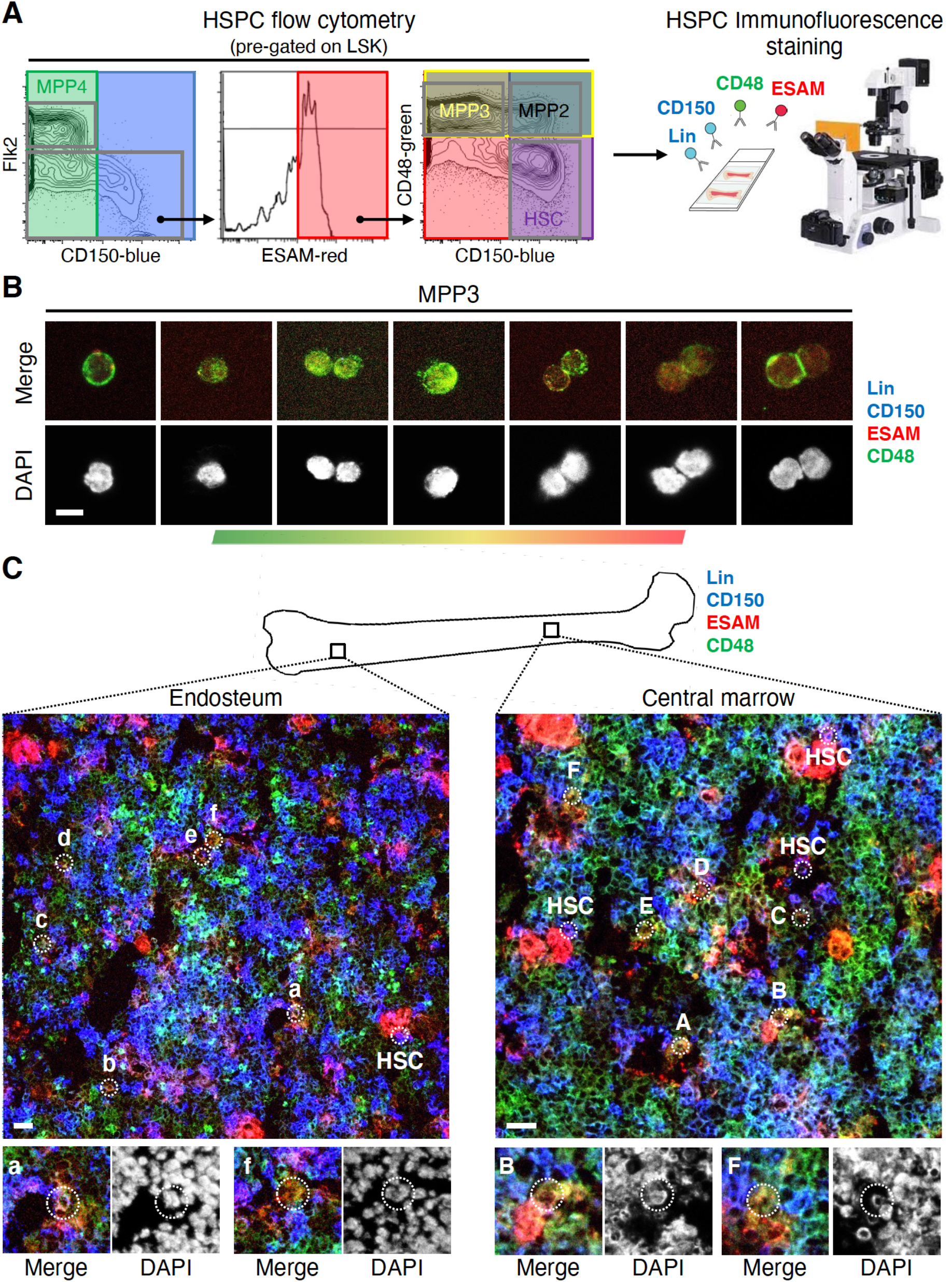
Localization of MPP3 in the vicinity of other HSPCs in the BM cavity. (**A**) Immunofluorescence staining strategy to identify hematopoietic stem and progenitor cell (HSPC) populations *in situ*. While HSC (purple), MPP2 (white), MPP3 (yellow) can be distinguished from the rest of the BM (blue) and megakaryocytes/megakaryocytic lineage (red) cells, MPP4 (green) are largely overlapping with GMPs in this staining scheme. (**B**) Representative images of isolated MPP3 stained with the immunofluorescence scheme for *in situ* imaging showing a gradient of yellow coloring. Scale bar, 10 μm. (**C**) Representative images of *in situ* immunofluorescence staining of HSPCs at the endosteum (left) and in the central marrow cavity (right). MPP3 are indicated by white dotted line circles, with magnified images of the indicated cells shown below with DAPI counterstain. HSCs are also denoted at both locations. Scale bar, 20 μm.

**Fig. S4.**
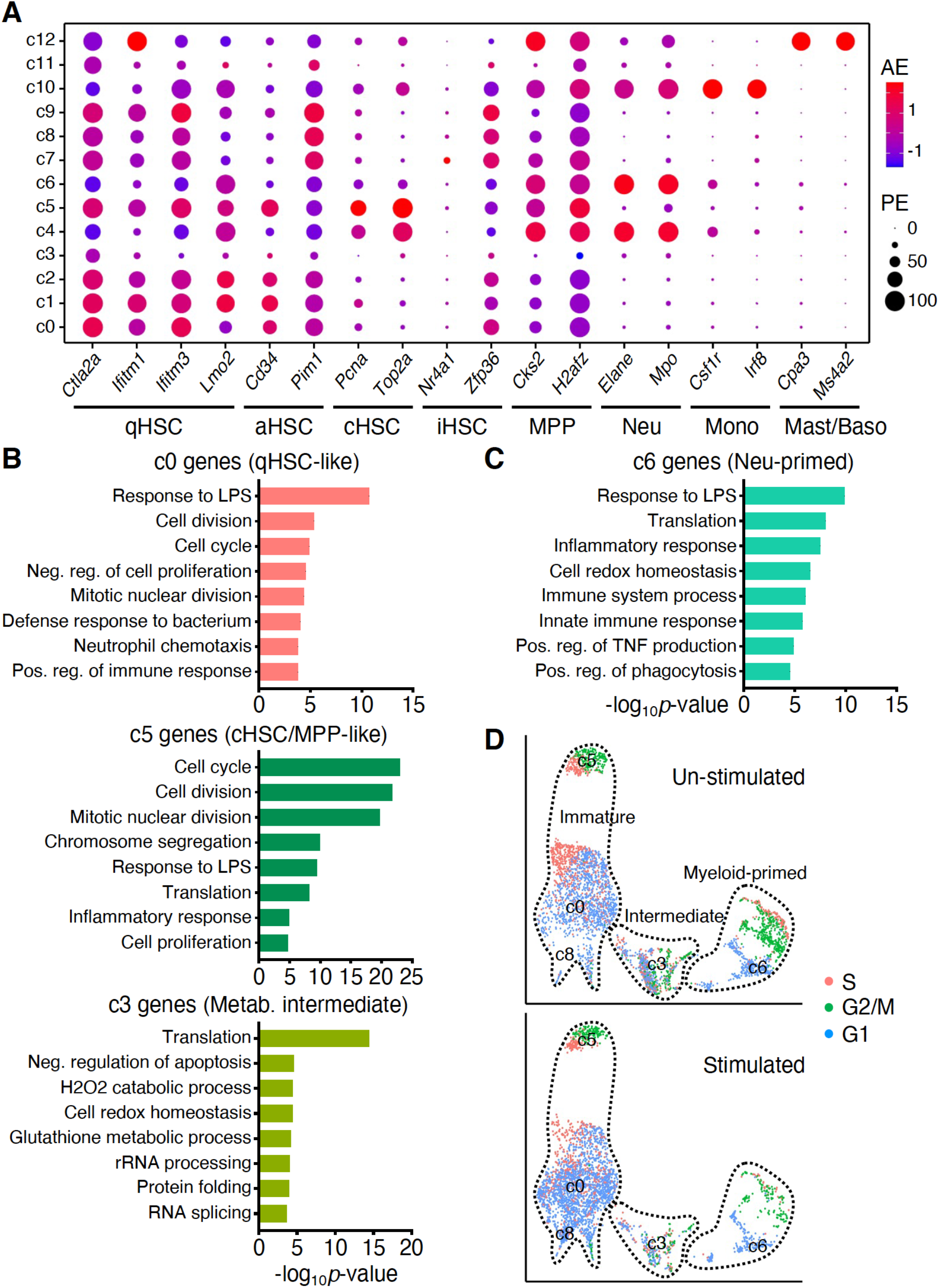
MPP3 heterogeneity. (**A**) Dot plot showing the expression of representative genes used for identifying clusters (c0 to c12) in the un-stimulated/stimulated MPP3 scRNAseq dataset. AE, average expression; PE, percent expressed; qHSC, quiescent HSC; aHSC, activated HSC; cHSC, cycling HSC; iHSC, inflammatory HSC; MPP, multipotent progenitor; Neu, neutrophil; Mono, monocyte; Mast, mast cell; Baso, basophil. (**B** and **C**) Gene ontology (GO) analyses for the indicated immature/intermediate (B) and myeloid-primed (C) clusters. The full list of GO analyses of all clusters is presented in Extended Data Table 2. (**D**) UMAP of un-stimulated/stimulated MPP3 showing cell cycle distribution. Un-stimulated/stimulated, 6 hours ± LPS/Pam3CSK4 (L/P) stimulation.

**Fig. S5.**
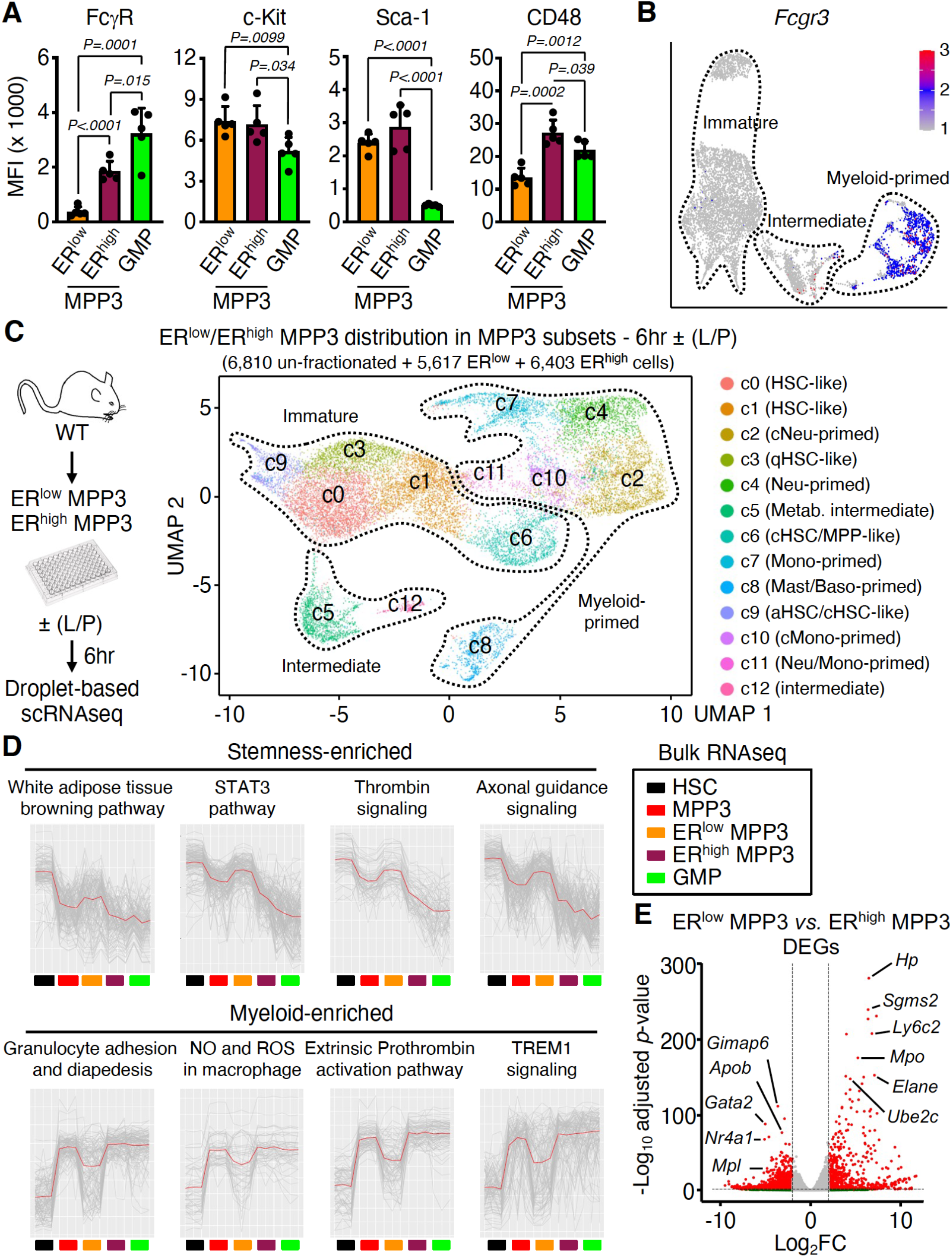
Molecular characterization of MPP3 subsets. (**A**) Quantification of surface marker expression in MPP3 subsets and GMPs. Results are shown as mean fluorescence intensity (MFI). Data are means ± S.D. and significance was assessed by a two-tailed unpaired Student’s t-test. (**B**) UMAP of un-stimulated/stimulated MPP3 showing *Fcgr3* expression. (**C**) UMAP of harmonized un-stimulated/stimulated ER^low^ MPP3, ER^high^ MPP3, and total MPP3 scRNAseq datasets with experimental scheme and cluster identification; ± (L/P), 6 hours *in vitro* treatment with or without LPS/Pam3CSK4; qHSC; quiescent HSC; aHSC, activated HSC; cHSC, cycling HSC; cNeu, cycling neutrophil; Neu, neutrophil; cMono, cycling monocyte; Mono, monocyte; Baso, basophil, Mast, mast cell; Metab., metabolic intermediate. (**D**) K-means clustering analysis of highly variable genes in HSC, MPP3, ER^low^ MPP3, ER^high^ MPP3 and GMP bulk RNAseq dataset showing representative enriched pathways. (**E**) Volcano plot of differentially expressed genes (DEG) between ER^low^ MPP3 vs. ER^high^ MPP3 showing representative examples. The full list of DEGs is presented in Extended Data Table 3.

**Fig. S6.**
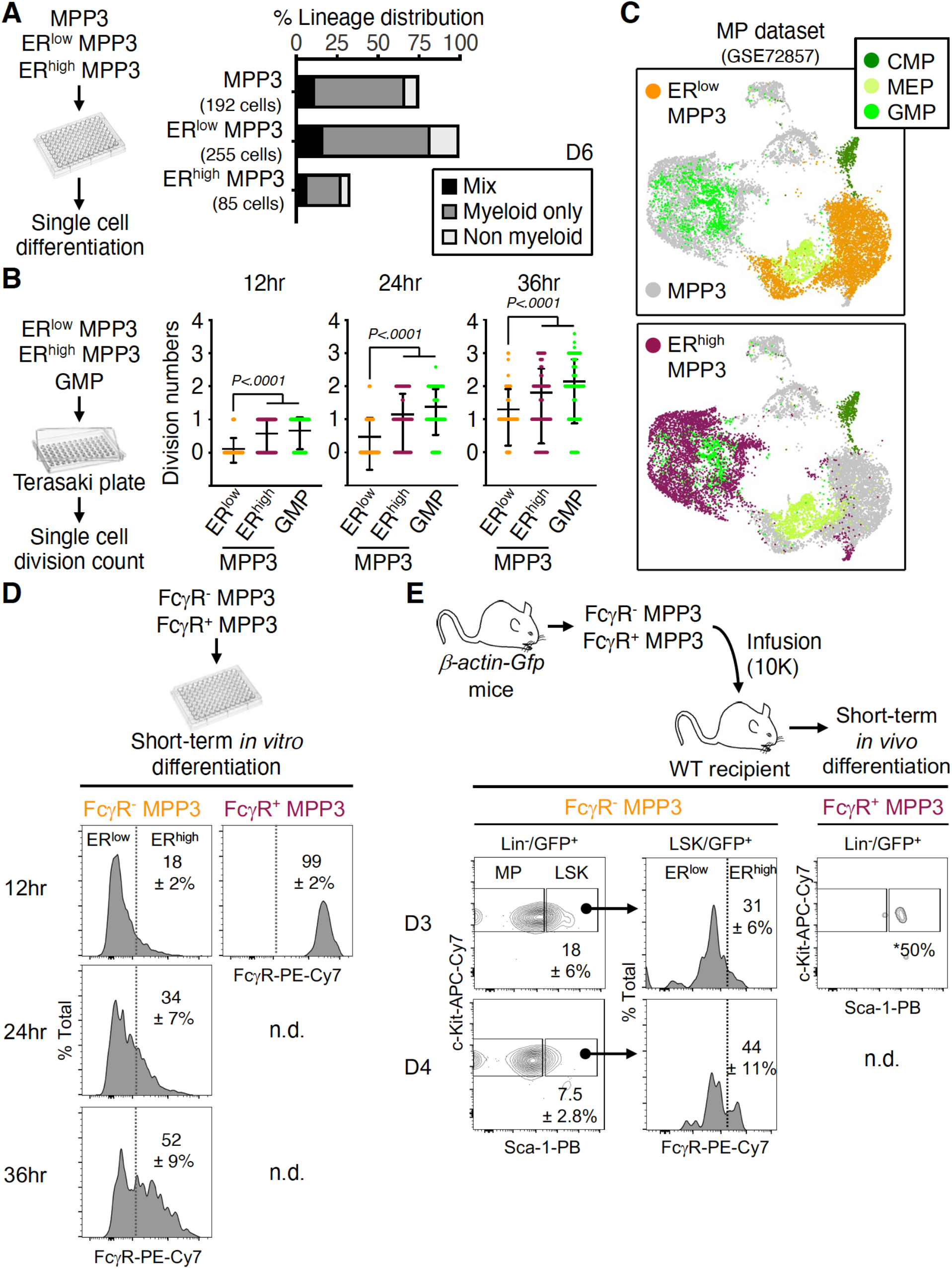
Functional characterization of MPP3 subsets. (**A**) Single cell *in vitro* lineage differentiation assay of MPP3 and ER^high^/ER^low^ MPP3 subsets with flow cytometry identification of single-cell derived colony composition after 6 days (D6) in culture. A total of 256 single cells were assessed in 3 independent experiments with data expressed as percentage of mix, myeloid only and non-myeloid lineage output. Myeloid, CD45^+^/Mac-1^+^/Gr-1^+^ cells; megakaryocyte, CD45^+^/Mac-1^-^/Gr-1^-^/CD41^+^/CD61^+^ cells and small manually counted megakaryocyte colonies; erythroid, CD45^+^/Mac-1^-^/Gr-1^-^/CD41^-^/CD61^+^/CD71^+^ cells; mast, CD45^+^/Mac-1^-^/Gr-1^-^/CD41^-^/CD71^-^/FcεRI^+^ cells; myeloid only, just myeloid output; non-myeloid, every other output but myeloid; mix, both myeloid and non-myeloid output. (**B**) Single cell *in vitro* division assay of ER^high^/ER^low^ MPP3 subsets and GMPs in Terazaki plates with assessment of cell division after 12 to 36 hours in culture. A total of 160 single cells were assessed in 3 independent experiments with data expressed as scatter dot plot (bar, mean). (**C**) UMAP of harmonized scRNAseq datasets projecting total MPP3 and ER^high^/ER^low^ MPP3 subsets with the published myeloid progenitor (MP) dataset (GSE72857); CMP, common myeloid progenitor; MEP, megakaryocyte/erythrocyte progenitor. (**D**) Short-term *in vitro* differentiation of FcγR^-^ and FcγR^+^ MPP3 subsets after 12 to 36 hours in culture (n =3). Representative FACS plots and quantification of FcγR^high^ frequencies are shown; n.d., not detected. (**E**) Short-term *in vivo* differentiation of FcγR^-^ and FcγR^+^ MPP3 subsets. Donor cells were isolated from *β-actin-Gfp* mice and infused into wild type (WT) recipients (10,000 cells per mouse, 3 to 4 recipients/population). GFP^+^ donor-derived cells were analyzed for contribution to the LSK (Lin^-^/Sca-1^+^/c-Kit^+^) BM compartment and FcγR expression at 3 and 4 days (D) post-infusion. Representative FACS plots and quantification of donor-derived LSK and FcγR^high^ frequencies are shown. *, donor-derived contribution was only detected in one of the 4 FcγR^+^ MPP3-infused recipients analyzed at day 3. Data are means ± S.D., and significance was assessed by a two-tailed unpaired Student’s t-test.

**Fig. S7.**
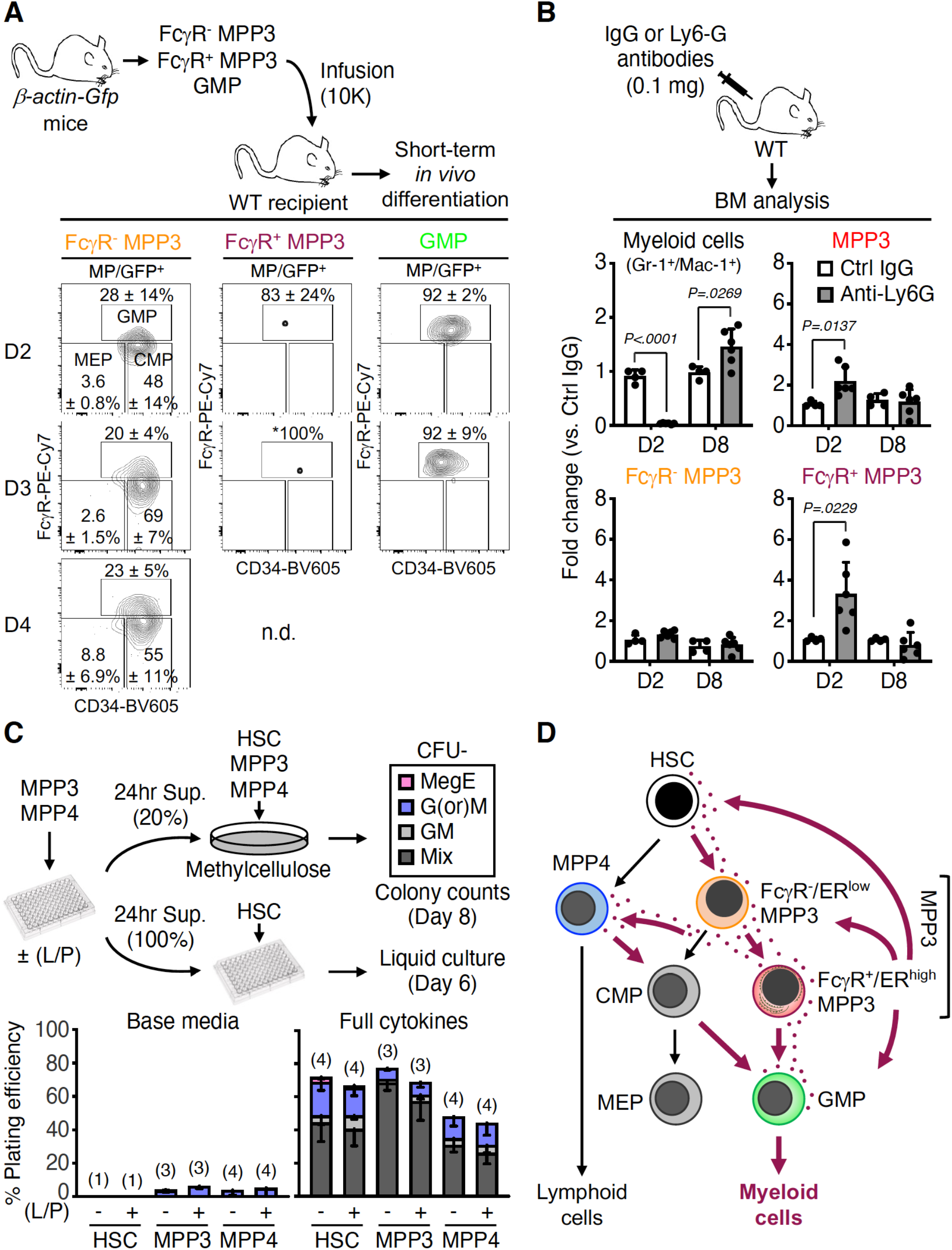
Characterization of FcγR^+^/ER^high^ MPP3 as a short-lived transitional population toward GMP commitment. (**A**) Short-term *in vivo* differentiation of FcγR^-^ and FcγR^+^ MPP3 subsets and GMPs. Donor cells were isolated from *β-actin-Gfp* mice and infused into wild type (WT) recipients (10,000 cells per mouse; 3 to 4 recipients/population). GFP^+^ donor-derived cells were analyzed for contribution to the myeloid progenitor (MP, Lin^-^/Sca-1^-^/c-Kit^+^) BM compartment at 2, 3 and 4 days (D) post-infusion. Representative FACS plots and quantification of donor-derived CMP, GMP and MEP frequencies are shown; n.d.; not detected. *, donor-derived contribution was only detected in one of the 4 FcγR^+^ MPP3-infused recipients analyzed at day 3. (**B**) Expansion of FcγR^-^ and FcγR^+^ MPP3 subsets during myeloid regeneration. WT mice were injected with control IgG or anti-Ly6G depleting antibodies and analyzed for changes in the indicated BM populations after 2 and 8 days (D). Results are expressed as fold changes in population size at each time point compared to IgG-treated mice. (**C**) Experimental scheme to assess the pro-myeloid differentiation effect of MPP3 and MPP4 supernatants on naïve HSCs, MPP3 and MPP4 plated either in methylcellulose with 20% supernatant or liquid cultures in 100% supernatant. Supernatants were collected upon culture of 10,000 MPP3 or MPP4 for 24 hours in 150 µl base media with or without (±) LPS/Pam3CSK4 (L/P) stimulation. Colony-forming units (CFU) in methylcellulose assays were scored after 8 days, and differentiating cells in liquid cultures were analyzed by flow cytometry after 6 days. Results from control methylcellulose assays performed with only base media or base media with full cytokine cocktail are shown in the bottom graphs; Mix, mixture of all lineages; GM, granulocyte/macrophage; G(or)M, granulocyte or macrophage; MegE, megakaryocyte/ erythrocyte. (**D**) Model depicting myeloid differentiation trajectories in early hematopoietic hierarchy and the role of secretory FcγR^+^/ER^high^ MPP3 subset in amplifying myeloid cell production. Data are means ± S.D., and significance was assessed by a two-tailed unpaired Student’s t-test.

**Fig. S8.**
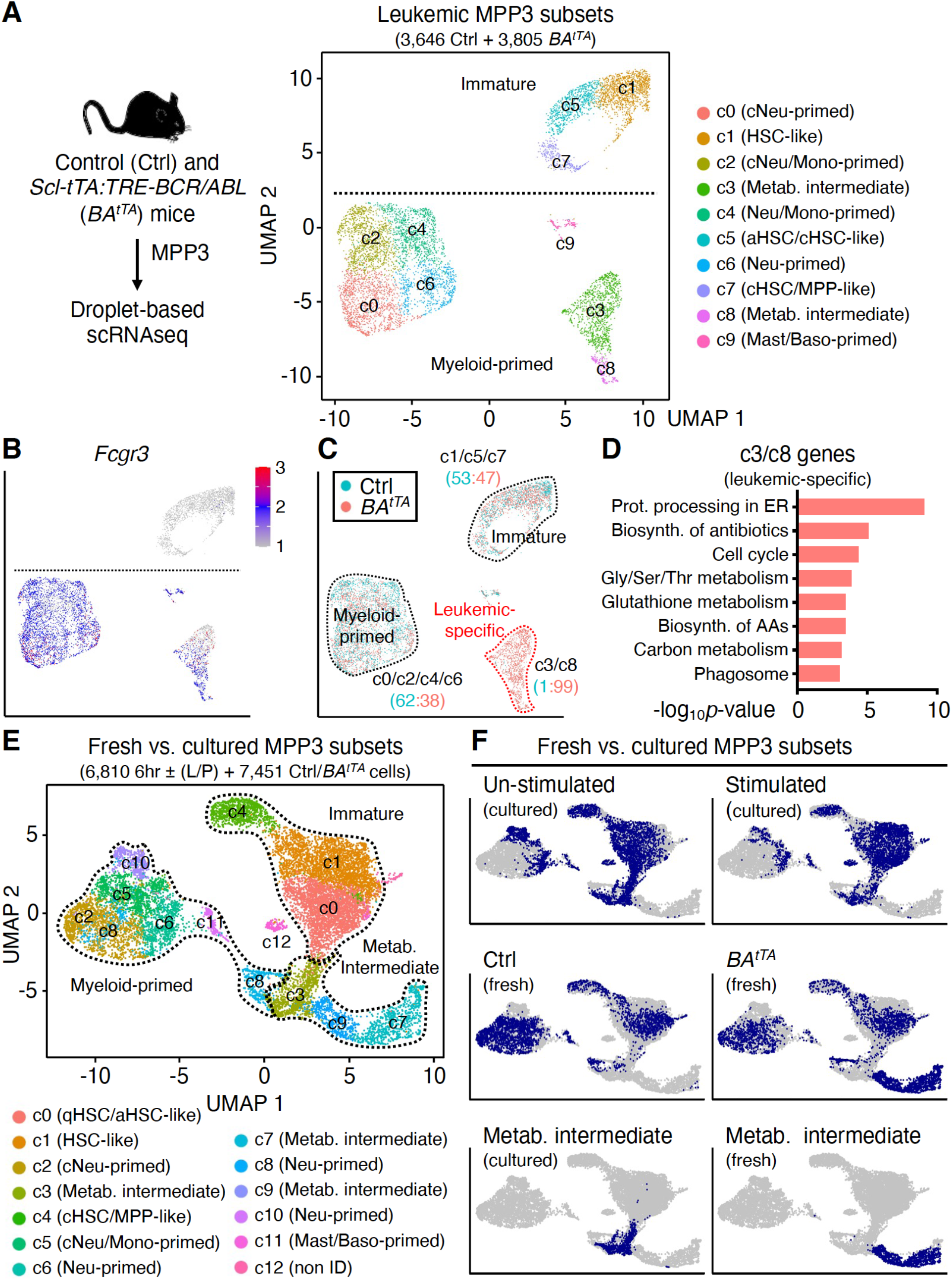
Molecular rewiring of leukemic MPP3. (**A**) UMAP representation of control (Ctrl) and *BA^tTA^* MPP3 scRNAseq dataset with experimental scheme and cluster identification; aHSC, activated HSC; cHSC, cycling HSC; cNeu, cycling neutrophil; Neu, neutrophil; Mono, monocyte; Baso, basophil, Mast, mast cell; Metab., metabolic intermediate. (**B** and **C**) UMAP representation of Ctrl/*BA^tTA^* MPP3 scRNAseq dataset showing (B) *Fcgr3* expression and (C) color coded ratio of Ctrl:*BA^tTA^* cells in major immature, myeloid-primed and leukemic-specific clusters. (**D**) KEGG pathway analysis of leukemic-specific (c3/c8) cluster genes; Prot, protein; Gly, glycine; Ser, serine; Thr, threonine; AAs; amino acids. (**E** and **F**) Comparison of freshly isolated and cultured MPP3 with: (e) UMAP of harmonized un-stimulated/stimulated MPP3 and Ctrl/*BA^tTA^* MPP3 scRNAseq datasets with cluster identification (± (L/P), 6 hours *in vitro* treatment with or without LPS/Pam3CSK4; qHSC; quiescent HSC; non ID, not identified), and (f) single projection of each dataset and specific metabolic (Metab.) intermediate clusters.

**Fig. S9.**
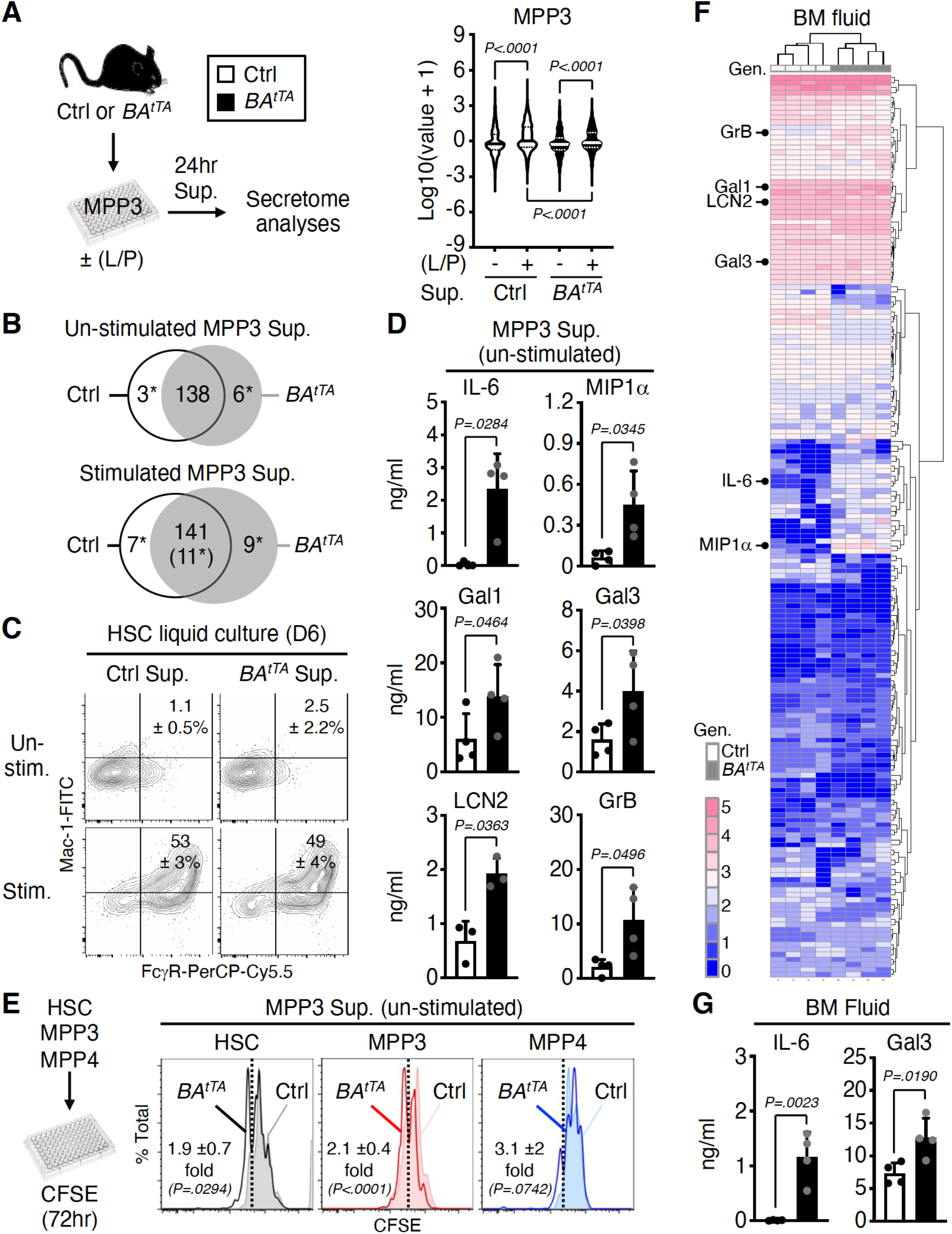
Constitutive cytokine secretion in leukemic MPP3. (**A** and **B**) Secretome analyses of un-stimulated and stimulated control (Ctrl) and *BA^tTA^* MPP3 supernatants (Sup.) with: (A) experimental scheme and violin plots of secreted cytokine intensity (n = 4; bar, median; dotted line, quartiles), and (B) Venn diagrams showing similarly and differentially secreted cytokines (*, significant change). Supernatants were collected upon culture of 10,000 Ctrl or *BA^tTA^* MPP3 for 24 hours in 150 µl base media with or without (±) LPS/Pam3CSK4 (L/P) stimulation. The full list of differentially secreted cytokines is provided in Extended Data Table 4. (**C**) Differentiation of naïve HSCs in Ctrl and *BA^tTA^* MPP3 supernatants analyzed after 6 days (D) of liquid culture for myeloid cell markers. Representative FACS plots and quantification of Mac-1^+^/FcγR^+^ frequencies are shown (n = 3). (**D**) Quantification of IL-6, MIP1a, Galectin 1 (Gal1), Galectin 3 (Gal3), Lipocalin 2 (LCN2) and Granzyme B (GrB) levels in un-stimulated Ctrl and *BA^tTA^* MPP3 supernatants. Significance was assessed by a two-tailed paired Student’s t-test. (**E**) Effect of un- stimulated Ctrl and *BA^tTA^* MPP3 supernatants on naïve HSCs, MPP3 and MPP4 proliferation analyzed by CFSE dilution assay after 72 hours in culture. Experimental scheme and representative FACS plots are shown. Dotted lines identify CSFE^low^ fast proliferative cells, and results indicate the pro-proliferative effect of *BA^tTA^* MPP3 supernatant shown as fold change compare to Ctrl MPP3 supernatant (n = 4-6). (**F** and **G**) Analysis of the cytokines secreted in the bone marrow (BM) fluid of Ctrl and *BA^tTA^* mice with: (F) heatmap of unsupervised clustering of BM fluid cytokine levels after quantile normalization (Gen; genotype), and (G) detailed quantification of IL-6 and Gal3 levels. BM fluids were obtained by flushing 4 long bones (femur and tibia) of each mouse with 200 µl media and were analyzed with the Raybiotech 200 mouse cytokine array. The 6 cytokines constitutively secreted by *BA^tTA^* MPP3 are indicated on the left in (F), and the full list of the cytokines differentially expressed in *BA^tTA^* vs. Ctrl BM fluids is provided in Extended Data Table 5. Data are means ± S.D. except when indicated, and significance was assessed by a two-tailed unpaired Student’s t-test except when indicated.

**Fig. S10.**
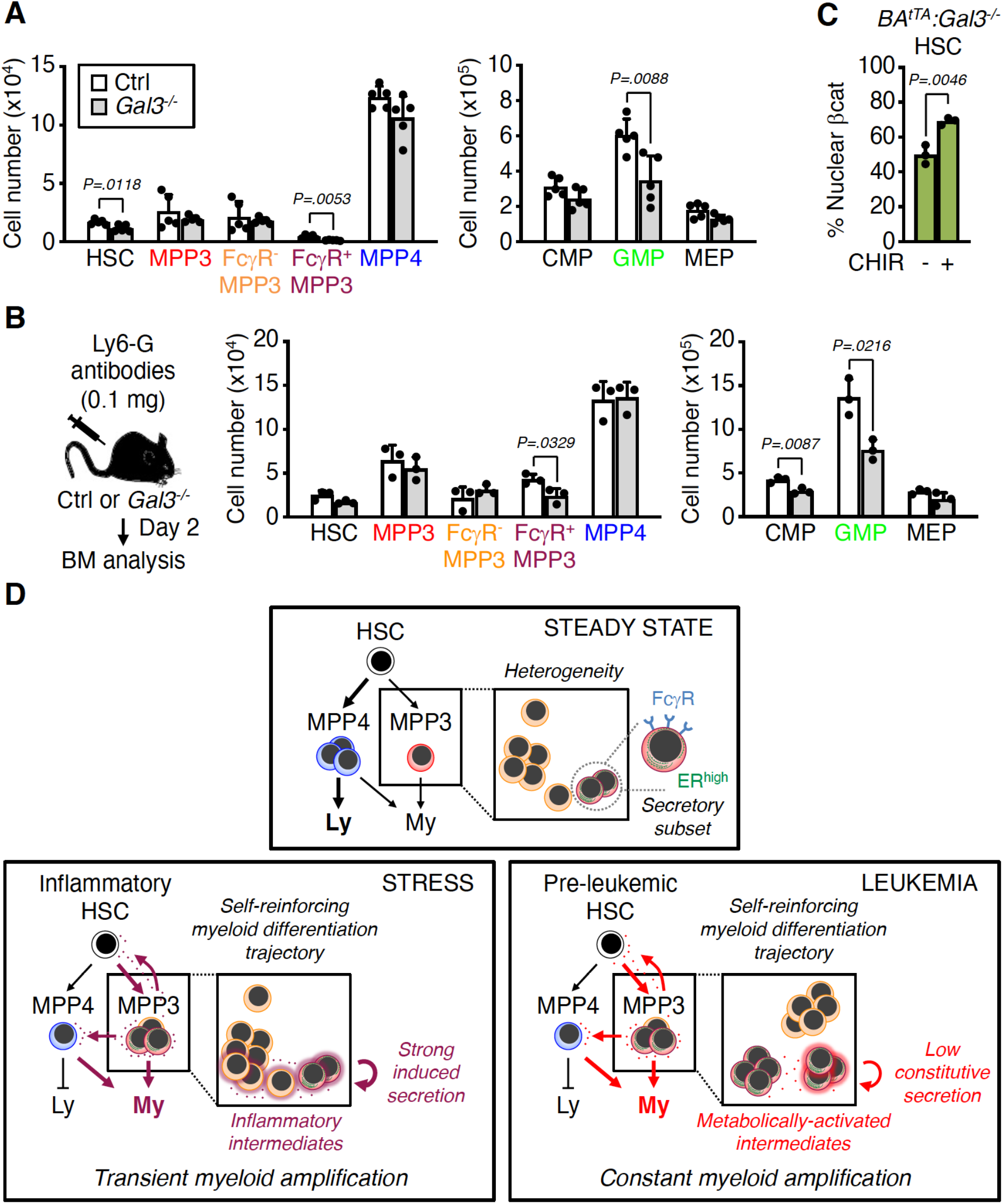
Importance of Galectin 3 secretion for myeloid amplification by leukemic MPP3. (**A**) HSPC and myeloid progenitor population size in age-matched control (Ctrl) *Gal3^+/+^* and knockout *Gal3^-/-^* mice. (**B**) Changes in HSPC and myeloid progenitor population size in Ctrl and *Gal3^-/-^* mice 2 days (D) after injection of anti-Ly6G depleting antibodies. (**C**) Changes in the frequency of nuclear β-catenin (βcat) positive *BA^tTA^:Gal3^-/-^* HSCs upon 18 hours *in vitro* treatment with the GSK3β inhibitor CHIR 99021 (CHIR, 30 µM). (**D**) Model depicting the novel regulatory function of a subset of secretory FcγR^+^/ER^high^ MPP3 that controls myeloid differentiation through lineage-priming and cytokine production. Although FcγR^+^/ER^high^ MPP3 act as a self-reinforcing compartment amplifying myeloid cell production in both stress and disease conditions, the actual mechanisms of lineage priming and secretion types are context dependent and specific to inflammatory signals or leukemia-driven genetics events. Data are means ± S.D., and significance was assessed by a two-tailed unpaired Student’s t-test.

**Table S1.** **Representative HSPC secretome clusters.** List of cytokines showing decreased (down), increased (up), or constitutively high (high) secretion upon stimulation in HSC, MPP4 and GMP supernatants. Supernatants were collected upon culture of 10,000 cells for 24 hours in 150 µl base media with or without LPS/Pam3CSK4 stimulation.

| HSC |  | MPP4 |  | GMP |  |
| --- | --- | --- | --- | --- | --- |
| **C1**<br>(down) | Fcg RIIB<br>Endoglin<br>VEGF R3<br>IL-1 R4<br>IL-12p70<br>JAM-A | **C4**<br>(down) | IL-23<br>TWEAK R<br>Nope<br>OPN<br>MIP-1γ<br>HGF<br>Lymphotactin<br>sFRP-3 | **C6**<br>(down) | Galectin-3<br>TWEAK R<br>Nope<br>MIP-1γ<br>Galectin-1<br>HGF<br>Lymphotactin<br>sFRP-3 |
| **C2**<br>(high) | Nope<br>OPN<br>TWEAK R | **C5**<br>(up) | MIP-1α<br>GM-CSF<br>TCA-3<br>OPG<br>CXCL16<br>IL-1α<br>MDC<br>MIP-2<br>RANTES<br>ICAM-1<br>IL-6<br>MIP-1β | **C7**<br>(down) | BLC<br>L-Selectin<br>IGF-1<br>CD40L<br>VEGF<br>Fcg RIIB<br>Endoglin<br>VEGF R1<br>ALK-1<br>P-Cadherin<br>TNF RI<br>IL-12p70<br>Gas 6 |
| **C3**<br>(up) | MIP-2<br>NOV<br>MIP-1γ<br>RANTES<br>ICAM-1<br>IL-6<br>MIP-1α<br>OPG<br>MIP-1β | | | **C8**<br>(up) | Prolactin<br>GITR<br>VCAM-1<br>IL-1α<br>MDC<br>Pentraxin 3<br>GM-CSF<br>MIP-1β<br>MCP-1<br>IL-1ra<br>TNFα<br>Progranulin |
| | | | | **C9**<br>(up) | IL-6<br>MIP-1α<br>Lipocalin-2<br>G-CSF<br>KC<br>MIP-2<br>OPN<br>TNF RII |

**Table S2. (separate file) Gene ontology (GO) analyses of MPP3 scRNAseq clusters and list of cluster-defining genes.** GO term and cluster-defining genes are shown for each cluster (c0 to c12); count, number of genes contributing to the GO term; %, percentage of cluster-defining genes contributing to the GO term.

**Table S3. (separate file) Differentially expressed genes (DEGs) between ER^high^ and ER^low^ MPP3.** List of significantly differentially expressed genes (> 4-fold change and p-value (FDR adjust) < 0.05) between ER^high^ MPP3 vs. ER^low^ MPP3.

**Table S4.** Differentially secreted cytokines in Ctrl and BA^tTA^ MPP3 supernatants. List of cytokines showing significantly higher (up) or lower (down) secretion in supernatants (Sup.) from control (Ctrl) or *BA^tTA^* MPP3 without (left) or with (right) 24 hours LPS/Pam3CSK4 stimulation. Cytokine levels were measured using the Raybiotech 200 mouse cytokine array (n = 4). Significance was assessed by a two-tailed paired Student’s t-test.

| Un-stimulated |  |  | Stimulated |  |  |  |
| --- | --- | --- | --- | --- | --- | --- |
|  | Ctrl Sup. | *BA<sup>tTA</sup>* Sup. |  | Ctrl Sup. | Overlap | *BA<sup>tTA</sup>* Sup. |
| Up | Tryptase E<br>VCAM1<br>IL-2 Ra | IL-6<br>MIP1α<br>Galectin 1<br>Galectin 3<br>Lipocalin 2<br>Granzyme B | Up | VEGF R2<br>MDC<br>VEGF<br>OPG<br>Cystatin C | MIP1β<br>RANTES<br>MIP2<br>MIP1γ<br>Lipocalin 2<br>MIP1α<br>TIM-1<br>GM-CSF<br>TNF R II<br>IL-6<br>OPN | KC<br>IL-1ra<br>Galectin 1<br>TNF RI<br>Gas 6<br>VCAM1<br>Fractalkine<br>G-CSF |
|  |  |  | Down | Kremen 1<br>Galectin 3 |  | Nope |

**Table S5.**
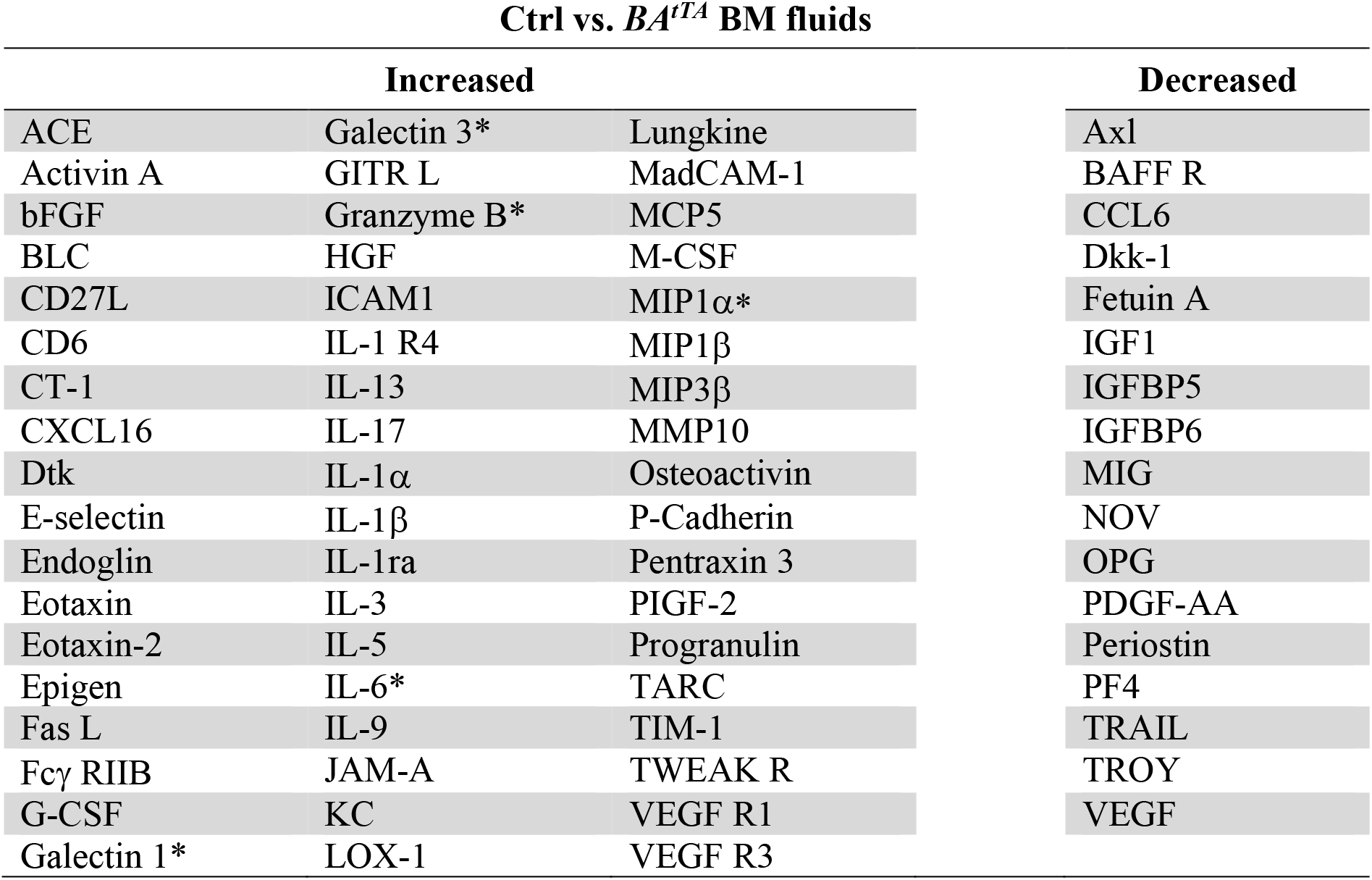
Differentially expressed cytokines in *BA^tTA^* BM fluids. List of cytokines showing increased (left) or decreased (right) expression in the bone marrow (BM) fluids from *BA^tTA^* mice compared to age-matched control (Ctrl) mice. Cytokine levels were measured using the Raybiotech 200 mouse cytokine array (n = 4) and significance was assessed by a two-tailed unpaired Student’s t-test. Asterisk (*) denotes cytokine constitutively secreted by *BA^tTA^* MPP3.

